# Early exposure to broadly neutralizing antibodies may trigger a dynamical switch from progressive disease to lasting control of SHIV infection

**DOI:** 10.1101/548727

**Authors:** Rajat Desikan, Rubesh Raja, Narendra M. Dixit

**Author notes:** Shared first authorship (Rajat Desikan, Rubesh Raja). Akamara Biomedicine Private Limited, Ist Floor, No. 465, Patparganj Industrial Area, Delhi, India, 110092 (Rajat Desikan).

## Abstract

Antiretroviral therapy (ART) for HIV-1 infection is life-long. Stopping therapy typically leads to the reignition of infection and progressive disease. In a major breakthrough, recent studies have shown that early initiation of ART can lead to sustained post-treatment control of viremia, raising hopes of long-term HIV-1 remission. ART, however, elicits post-treatment control in a small fraction of individuals treated. Strikingly, passive immunization with broadly neutralizing antibodies (bNAbs) of HIV-1 early in infection was found recently to elicit long-term control in a majority of SHIV-infected macaques, suggesting that HIV-1 remission may be more widely achievable. The mechanisms underlying the control elicited by bNAb therapy, however, remain unclear. Untreated infection typically leads to progressive disease. We hypothesized that viremic control represents an alternative but rarely realized outcome of the infection and that early bNAb therapy triggers a dynamical switch to this outcome. To test this hypothesis, we constructed a model of viral dynamics with bNAb therapy and applied it to analyse clinical data. The model fit quantitatively the complex longitudinal viral load data from macaques that achieved lasting control. The model predicted, consistently with our hypothesis, that the underlying system exhibited bistability, indicating two potential outcomes of infection. The first had high viremia, weak cytotoxic effector responses, and high effector exhaustion, marking progressive disease. The second had low viremia, strong effector responses, and low effector exhaustion, indicating lasting viremic control. Further, model predictions suggest that early bNAb therapy elicited lasting control via pleiotropic effects. bNAb therapy lowers viremia, which would also limit immune exhaustion. Simultaneously, it can improve effector stimulation via cross-presentation. Consequently, viremia may resurge post-therapy, but would encounter a primed effector population and eventually get controlled. ART suppresses viremia but does not enhance effector stimulation, explaining its limited ability to elicit post-treatment control relative to bNAb therapy.

**Author Summary:** In a remarkable advance in HIV cure research, a recent study showed that 3 weekly doses of HIV-1 broadly neutralizing antibodies (bNAbs) soon after infection kept viral levels controlled for years in most macaques treated. If translated to humans, this bNAb therapy may elicit a functional cure, or long-term remission, of HIV-1 infection, eliminating the need for life-long antiretroviral therapy (ART). How early bNAb therapy works remains unknown. Here, we elucidate the mechanism using mathematical modeling and analysis of *in vivo* data. We predict that early bNAb therapy suppresses viremia, which reduces exhaustion of cytotoxic effector cells, and enhances antigen uptake and effector stimulation. Collectively, these effects drive infection to lasting control. Model predictions based on these effects fit *in vivo* data quantitatively. ART controls viremia but does not improve effector stimulation, explaining its weaker ability to induce lasting control post-treatment. Our findings may help improve strategies for achieving functional cure of HIV-1 infection.

## Introduction

Current antiretroviral therapies (ART) for HIV-1 infection control viremia in infected individuals but are unable to eradicate the virus. ^1^ A reservoir of latently infected cells, which is established soon after infection^2^, escapes drugs and the host immune response^3^, is long-lived^4,5^, and can reignite infection following the cessation of therapy ^6^, presents the key obstacle to sterilizing cure. Efforts are now aimed at eliciting a “functional cure” of the infection, where the virus can be controlled without life-long treatment even though eradication is not possible. ^7^ That functional cure can be achieved has been demonstrated by the VISCONTI trial, where a subset of patients, following early initiation of ART, maintained undetectable viremia long after the cessation of treatment. ^8^ A limitation, however, is that the subset that achieves post-treatment control with ART is small, 5-15% of the patients treated. ^9^ In a major advance, Nishimura *et al.*^10^ found recently that early, short-term passive immunization with a combination of two HIV-1 broadly neutralizing antibodies (bNAbs) elicited lasting control of viremia in 10 of 13, or nearly 77%, of SHIV-infected macaques treated. This high success rate raises the prospect of achieving functional cure in all HIV-1 infected individuals using short-term bNAb therapy. Efforts have been initiated to develop immunotherapies that may further improve response rates in primate models^11–14^ and to translate them to humans ^15–17^.

Devising passive immunization protocols that would maximize the chances of achieving functional cure requires an understanding of the mechanism(s) with which early, short-term passive immunization with bNAbs induces sustained control of viremia. Nishimura *et al.*^10^ argue that the control they observed, lasting long (years) after the administered bNAbs were cleared from circulation (weeks), was due to the effector function of cytotoxic cells such as CD8^+^ T lymphocytes (CTLs) because transient suppression of effector cells, using anti-CD8 antibodies (Abs), well after the establishment of control resulted in a temporary surge of viremia. How short-term bNAb therapy leads to sustained effector stimulation, and therefore lasting viremic control, remains unknown. ^15^

Here, we propose a dynamical systems-based role of bNAbs in eliciting lasting control. We hypothesized that lasting viremic control is a distinct but rarely realized outcome of HIV-1 infection and that early, short-term bNAb therapy triggers a switch in disease dynamics that significantly increases the likelihood of realizing this outcome. To test this hypothesis and to elucidate the mechanisms with which bNAbs orchestrate this switch, we constructed a mathematical model of HIV dynamics under bNAb therapy and applied it to analyse recent *in vivo* data.

## Results

### Mathematical model of viral dynamics with passive immunization

We constructed a mathematical model that considered the within-host dynamics of populations of uninfected target CD4^+^ T cells, productively infected cells, free virions, effector cells, and administered bNAbs (Figure 1). The dynamics in the absence of bNAbs were similar to those in recent models of viral dynamics that include effector cells. ^18–20^ Briefly, uninfected target cells get infected by free virions to yield productively infected cells, which in turn produce more free virions. Effector cells are stimulated by infected cells and could become exhausted by sustained antigenic stimulation. Activated effectors kill infected cells. We modified these processes based on the known effects of bNAbs ^15,21^: bNAbs opsonize free virions, preventing them from infecting cells and enhancing their clearance. ^22–24^ Opsonized virus can also be taken up by immune cells, particularly macrophages, and via processes now being unraveled improve antigen presentation and increase infected cell death. ^25–28^ Thus, we let bNAbs 1) enhance the clearance of free virions and 2) increase effector activation, via enhanced antigen uptake, both in a dose-dependent manner. The enhanced viral clearance, which results in a reduction in viremia, was assumed to subsume the effect of virus neutralization (see Methods). By lowering antigen levels, bNAbs could reverse effector exhaustion.

**Fig. 1.**
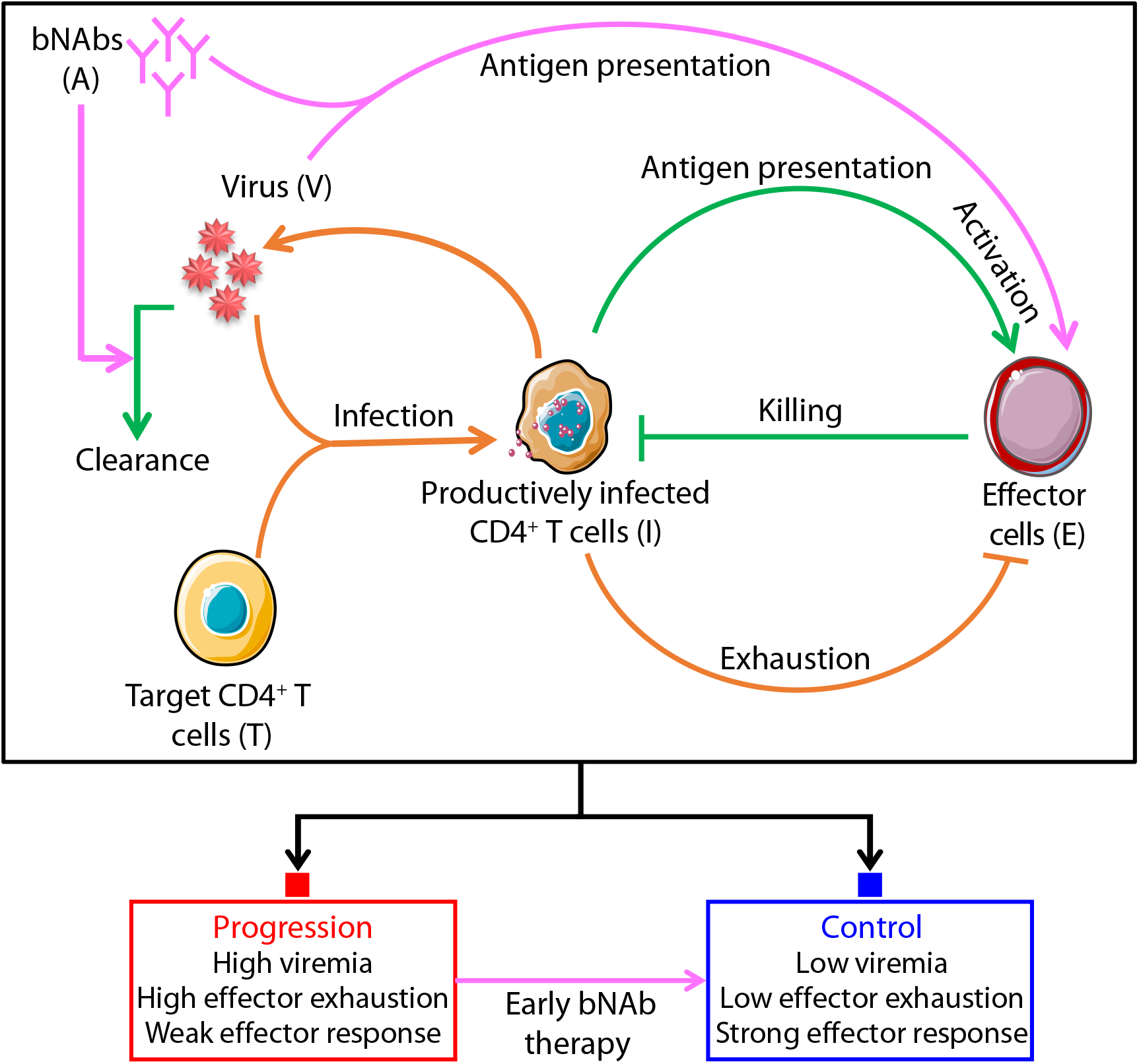
Schematic of the mathematical model of SHIV dynamics with bNAb therapy. Orange and green arrows indicate processes involved in the growth and control of infection, respectively, while magenta arrows indicate processes initiated or enhanced by bNAbs. Corresponding model equations are described in Methods. Steady state analysis of the model indicates two outcomes of the infection: chronic infection with high viremia that marks progressive disease (red filled square), typically realized in the absence of treatment, and viremic control (blue filled square), a switch to which is orchestrated by early bNAb therapy.

ART was assumed to reduce the productive infection of target cells. Anti-CD8 Abs were assumed to reduce the effector population and compromise host effector functions for a duration corresponding to the residence time of the depleting Abs in circulation.

We constructed equations to describe the resulting dynamics and solved them using parameter values representative of SHIV infection of macaques (Methods).

### Model predictions fit in vivo data

To test whether the model accurately captured the underlying dynamics of infection and the influence of bNAbs, we applied it to describe the viral load changes reported in Nishimura *et al.*^10^ where 3 weekly infusions of a combination of two HIV-1 bNAbs, 3BNC117 and 10-1074, starting from day 3 post SHIV challenge were administered. We considered the 10 “responder” macaques who showed no significant decline in CD4^+^ T cell counts and included 6 with undetectable post-treatment set-point viremia (< 10^2^ copies/mL), termed “controllers”, and 4 with detectable but low post-treatment set-point viremia (< 10^3^ copies/mL). We also considered 10 untreated macaques in the study with viral load measurements reported during the acute phase of infection. Where available, we fixed model parameter values from previous studies (Methods and Table 1). We estimated the remaining parameters by using non-linear mixed effects modelling to simultaneously fit viral load data (Figure 2) and bNAb concentrations (Figure 3) in the responder macaques and viral load data in the untreated macaques (Figure 4). The model provided good fits to the data, yielding population-level parameter estimates (Table 2) and parameter estimates yielding best-fits to individual macaques (Tables S1 and S2). The population parameters compared well with previous estimates, wherever available (Methods).

**Table 1.** Model parameters fixed from previous studies.

| Parameter | Meaning | Value | Units | Citation |
| --- | --- | --- | --- | --- |
| $d_T$ | Per capita death rate of uninfected CD4<sup>+</sup> T cells | 0.01 | day<sup>-1</sup> | Ref.<sup>18</sup> |
| $d_I$ | Per capita death rate of productively infected CD4<sup>+</sup> T cells | 0.5 | day<sup>-1</sup> | Ref.<sup>18</sup> |
| $c$ | Viral clearance rate | 38 | day<sup>-1</sup> | Refs.<sup>22,29,30</sup> |
| $k_E$ | Per capita proliferation rate of effectors | 0.1 | day<sup>-1</sup> | Refs.<sup>31,32</sup> |
| $q_c^*$ | Half-maximal Hill function threshold for exhaustion | 0.5 | - | Ref.<sup>19</sup> |
| $n$ | Hill coefficient for exhaustion | 4 | - | Ref.<sup>19</sup> |
| $d_q$ | Exhaustion reversal rate constant | 0.1 | day<sup>-1</sup> | Ref.<sup>19</sup> |
| $A_1^{dose}$ | Estimated dose of the bNAb 3BNC117 | 65000<sup>#</sup> | μg | Ref.<sup>10</sup> |
| $A_2^{dose}$ | Estimated dose of the bNAb 10-1074 | 65000<sup>#</sup> | μg | Ref.<sup>10</sup> |

<sup>#</sup> Nishimura *et al.* administer 10 mg/kg of bNAbs to the macaques, and we assume an average body weight of 6.5 kg.

**Table 2.**
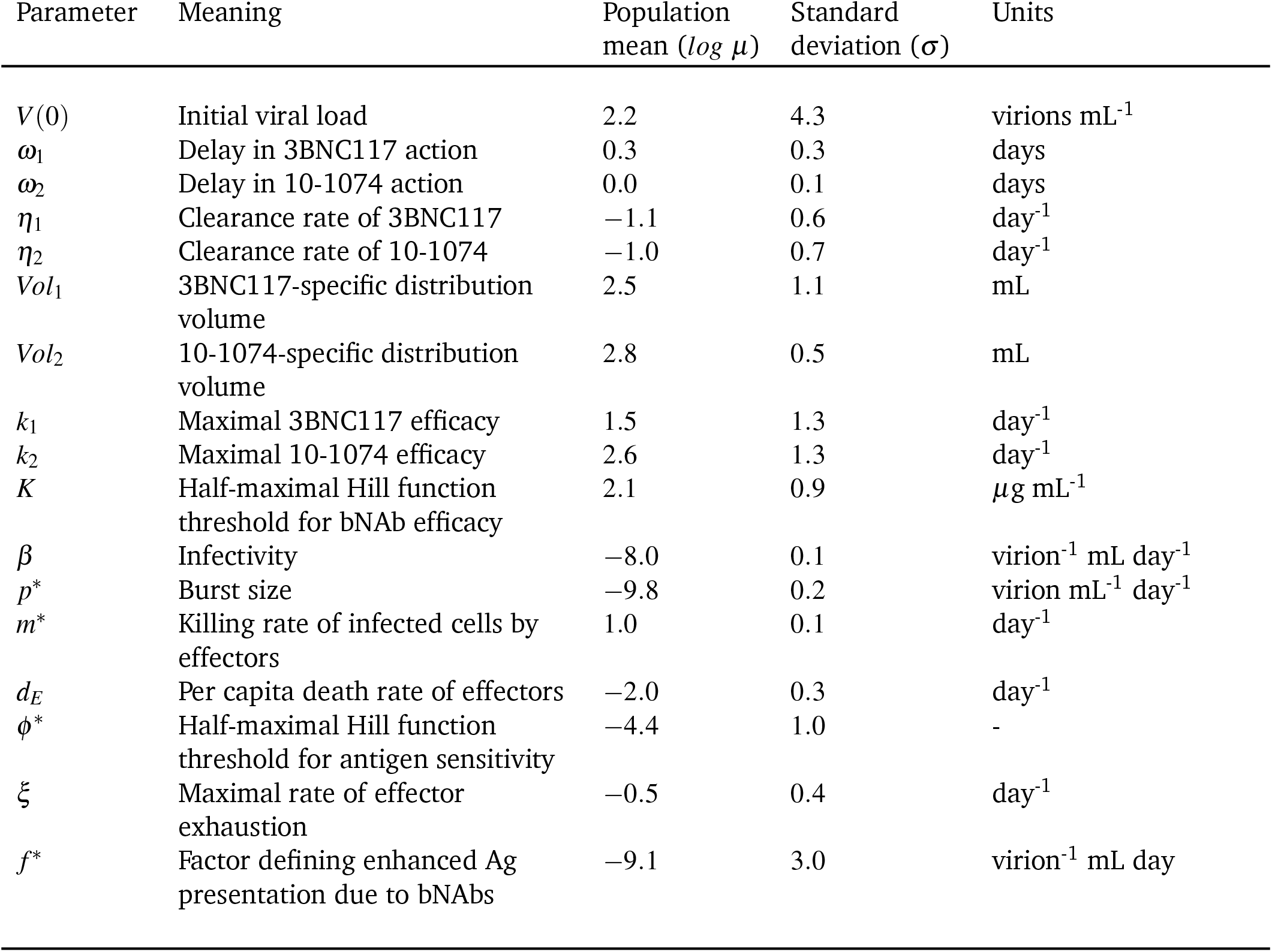
Mean population parameters and standard deviations estimated by simultaneously fitting our model (Eqs. 11–18) to data of *V*, *A*_1_ and *A*_2_ from untreated macaques and responders (see Methods).

**Fig. 2.**
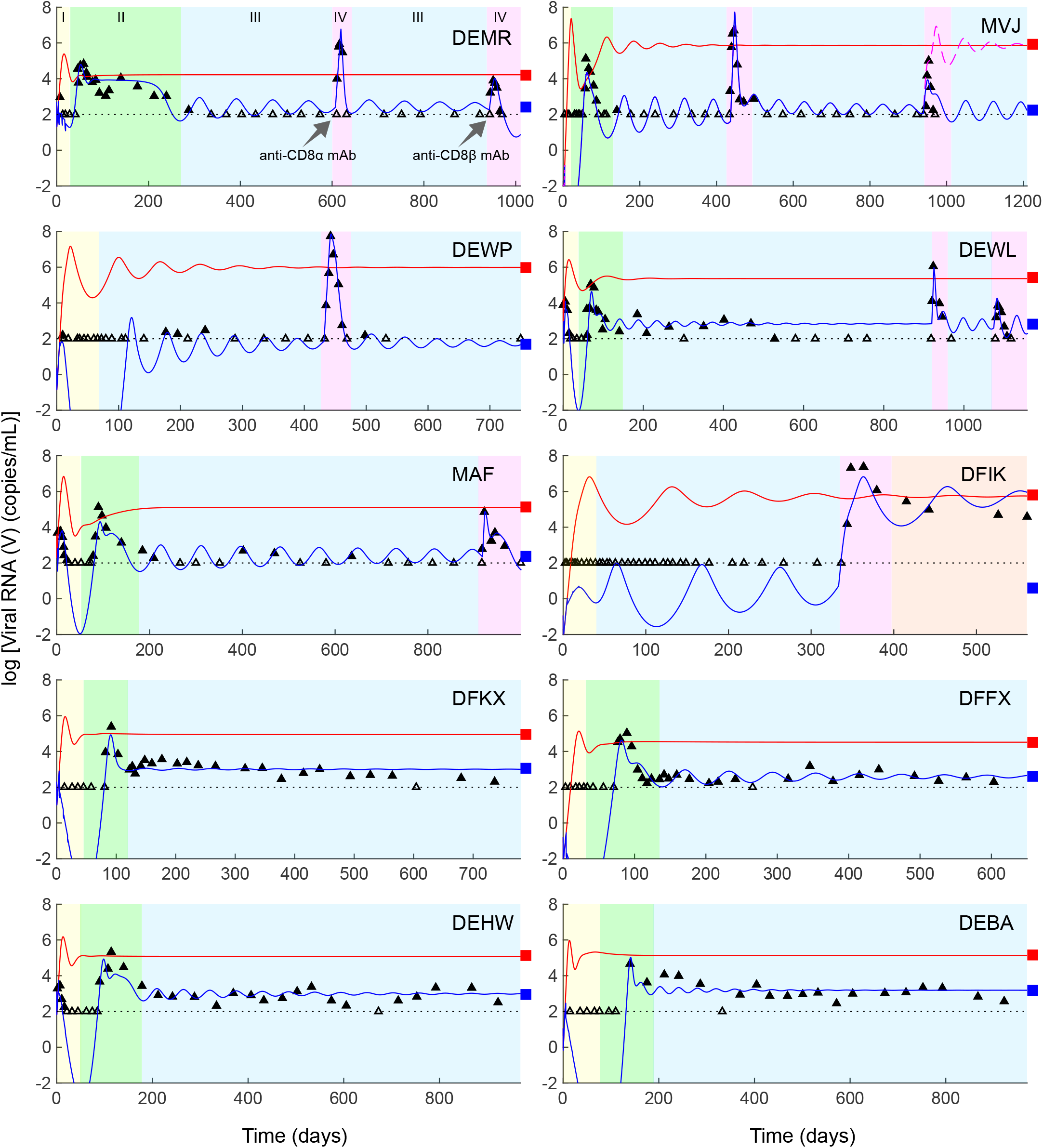
Model fits viral dynamics in responder macaques. Model fits (blue lines) to viral load data10 (symbols) from the 10 controller macaques, shown in individual panels. Empty symbols mark measurements showing undetectable viremia and filled symbols above detection. The latter were used for fitting and the former censored. Black dashed lines in all panels indicate the viral load detection limit (100 RNA copies/mL). The corresponding bNAb concentration dynamics are in Figure 3. In all panels, phase I (yellow) marks the duration when bNAbs are present in circulation, phase II (green) the viremic resurgence post the clearance of bNAbs, phase III (blue) the ensuing viremic control, and, where relevant, phase IV (pink), in two parts, the disruption of this control using anti-CD8*α* and anti-CD8*β* Abs, respectively. For the macaque MVJ, we demonstrate the loss of viral control that occurs when effector depletion levels are increased (magenta line), as is observed with the macaque DFIK. The digitized data used for the fitting is available as a supplementary excel file (Database.xlsx). In all cases, predictions without bNAb therapy are included for comparison (red lines). Parameter values used are in Tables 1 and S1. Macaques DEWP, MVJ, DFFX, DFKX, and DFIK were inoculated intrarectally and DEWL and MAF intravenously with 1000 TCID50 (50% tissue culture infective dose) of SHIVAD8-EO virus. DEMR, DEBA, and DEHW received 100 TCID_50_ intravenously.

**Fig. 3.**
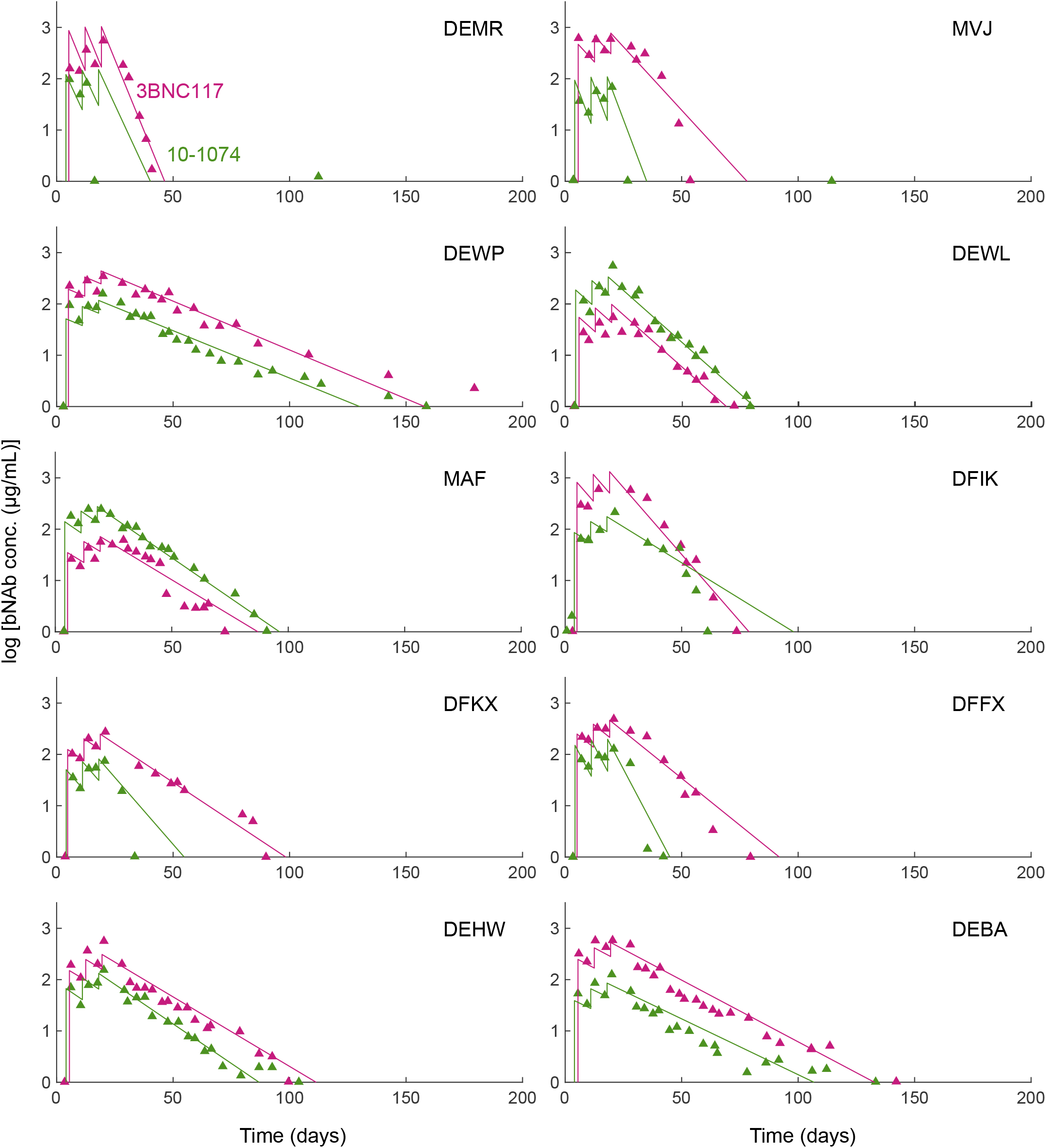
bNAb pharmacokinetics in responders. Fits of model predictions (lines) to data from individual macaques (symbols) of bNAb plasma concentrations for the ten responders in Nishimura *et al.* 10 obtained by simultaneously fitting our model (Eqs. 11–18) to *V*, *A*_1_ and *A*_2_ across all macaques. See Methods for the fitting procedure. The resulting parameter estimates are in Table S1. The bNAb serum half-lives averaged across all macaques are 9.2 days (range 2.7 − 15.8 days) for 3BNC117, and 8.4 days (range 2.5 − 16.3 days) for 10-1074.

**Fig. 4.**
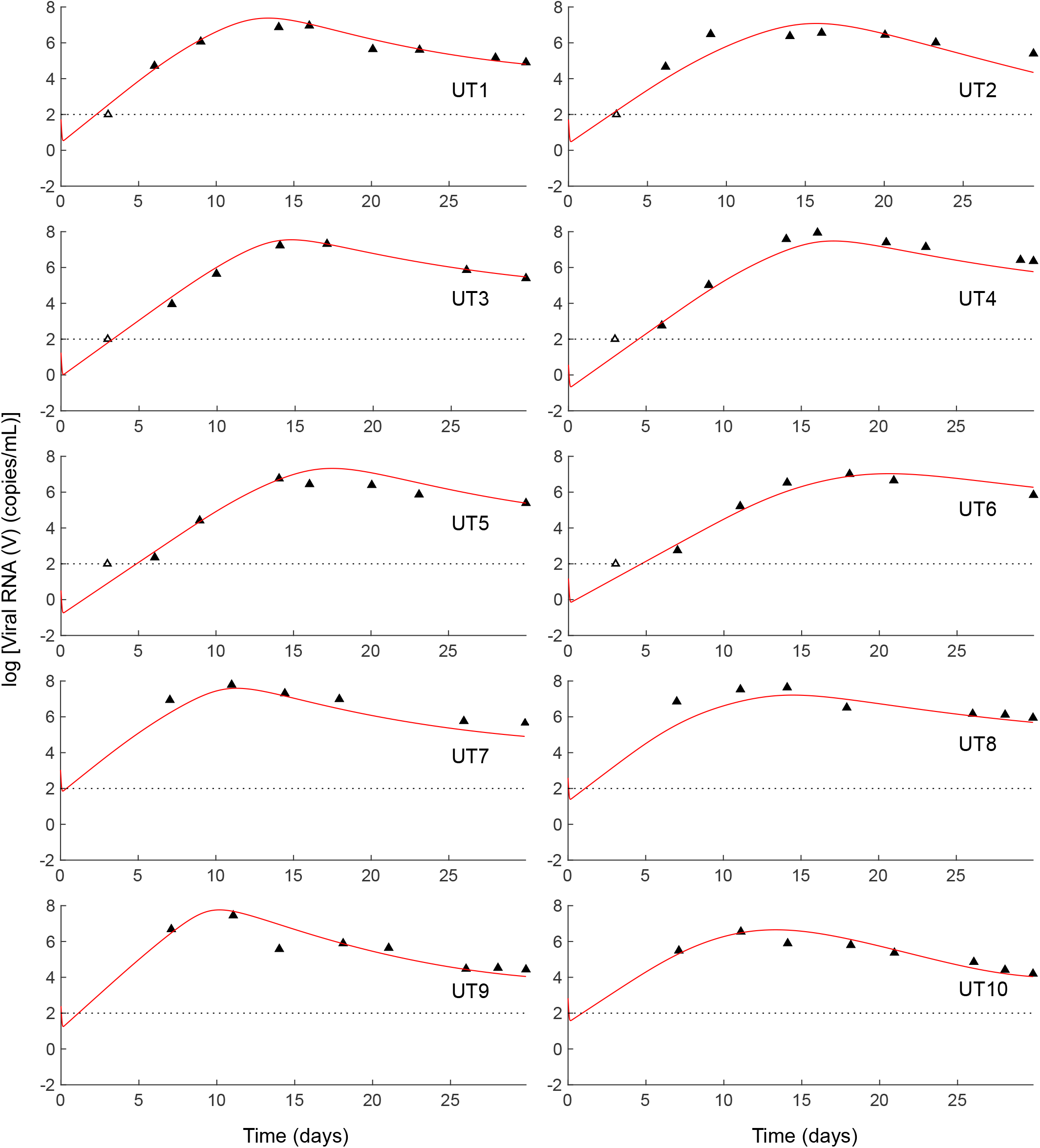
Untreated macaques. Fits (red lines) to the acute phase viral loads of the ten untreated macaques in Nishimura *et al.* ^10^ obtained by simultaneously fitting our model (Eqs. 11–18) to *V*, *A*_1_ and *A*_2_ across all macaques. See Methods for the fitting procedure. The best-fit parameter estimates are in Table S2.

The viral dynamics during and post bNAb therapy was complex and could be divided into the following phases (Figure 2). Viremia dropped post the acute infection peak to undetectable levels (< 10^2^ copies/mL), due to bNAb administration, where it remained until the administered bNAb level in circulation declined to a point where its ability to control viremia was lost (Phase I in Figure 2a). Viremia then resurged to an elevated level (~ 10^5^ copies/mL), well above the acute infection peak (< 10^4^ copies/mL), but subsequently declined spontaneously to a low (10^2^ − 10^3^ copies/mL) level (Phase II), where it remained for the rest of the duration of follow-up (>3 years; Phase III). The approach to steady state was a slowly dampened oscillation, similar to that seen in previous models of HIV-1 infection that considered an effector response^33–35^.

Following the administration of anti-CD8 Abs, several months after the establishment of control, viremia immediately resurged to high levels again, but was eventually controlled (Phase IV) in 5 of 6 macaques thus treated (Figure 2). One macaque (DFIK) lost control following the administration of anti-CD8 Abs and reached a high viral load set-point, akin to untreated macaques.

The ability to describe these complex viral load changes during and post bNAb therapy quantitatively gave us confidence in our model. We applied our model next to elucidate the mechanistic origins of the observed long-term viremic control.

### Viremic control is a distinct outcome of infection

We hypothesized, based on the two outcomes, progressive disease vs. viremic control, achieved following bNAb therapy as well as CD8 depletion, that lasting control observed with early bNAb therapy was a distinct steady state of the system. This hypothesis of bistability, or the existence of two distinct stable steady states, is consistent with a previous analysis of post-treatment control with ART ^18^ and may be a general feature of the interactions between antigen and effector cells ^20^. Indeed, solving our model equations (Methods), we identified two underlying stable steady states (Figure 5). The first had high viremia, relatively high levels of effector exhaustion and relatively weak effector function, marking chronic infection leading to progressive disease. The second had low viremia, relatively low levels of effector exhaustion and relatively strong effector function, indicating lasting control. These states existed over wide ranges of parameter values (Figure S1).

**Fig. 5.**
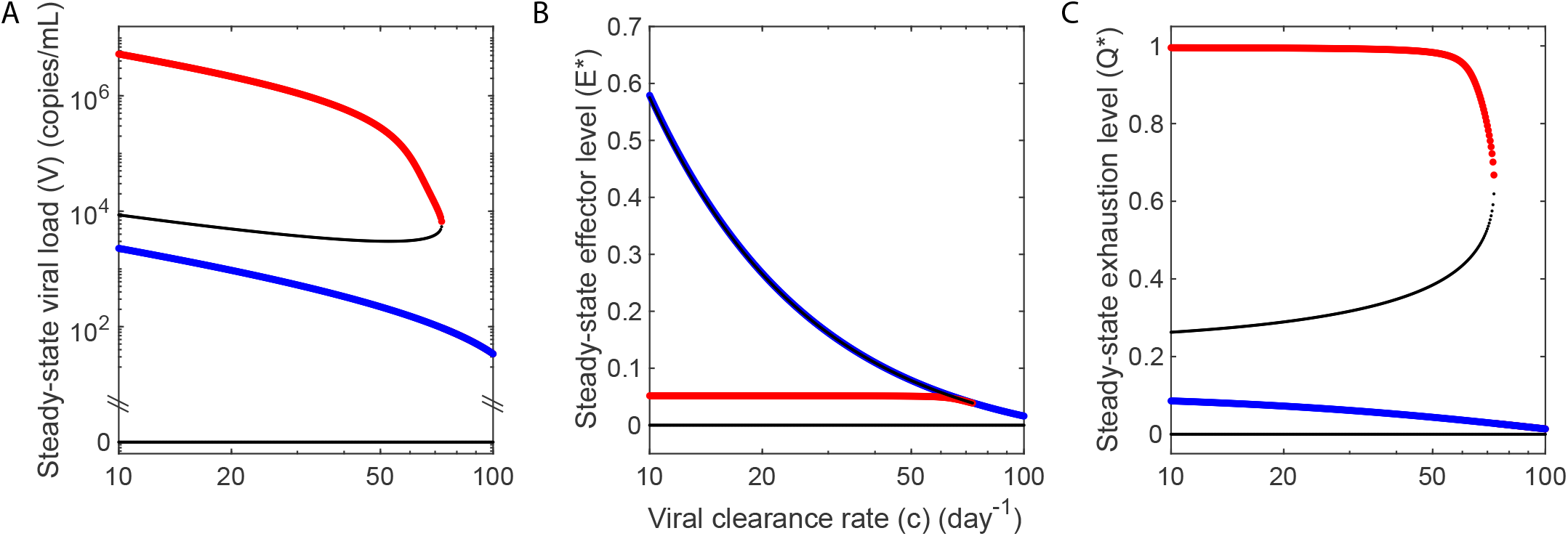
Bifurcation diagrams indicating the two distinct outcomes of progressive disease and viremic control. Model calculations of the steady state (**a**) viral load, (**b**) normalized effector level (*E*\* = *Ed*_*E*_/*ρ*), and (**c**) normalized level of exhaustion (*Q*\* = *Qd*_*q*_/*κ*), obtained by varying the viral clearance rate. The stable states of progressive disease and viremic control are shown in red and blue, respectively. Thin black lines represent unstable steady states. In (**b**), the intermediate unstable state lies close to the state of viremic control. The steady states are separated by other state variables, including the level of exhaustion, evident in (**c**). Parameter values used are in Tables 1 and 2. Bifurcation diagrams for other parameters are shown in Figure S1.

To indicate the two steady states for given parameter values, we use red and blue filled squares, respectively, in all the figures. The two stable states were separated by an unstable steady state of intermediate viremia, and intermediate effector stimulation and/or exhaustion levels. In addition, an uninfected steady state existed that was also unstable.

### Early, short-term bNAb therapy switches the outcome to viremic control via pleiotropic effects

When a new infection occurs, viremia rises significantly during the acute infection phase and achieves high levels typically before an adaptive effector response is mounted. ^36^ The effector response thus lags and is dominated by the virus. If left untreated, the high viremic state, leading to progressive disease, is realized, as observed with untreated macaques challenged with SHIV ^37,38^. Our model predictions are consistent with these observations (Figure 4; red lines in Figures 2 and 6). They show that following the initial transients, a high viremia, a high level of exhaustion, and a weak effector response result.

**Fig. 6.**
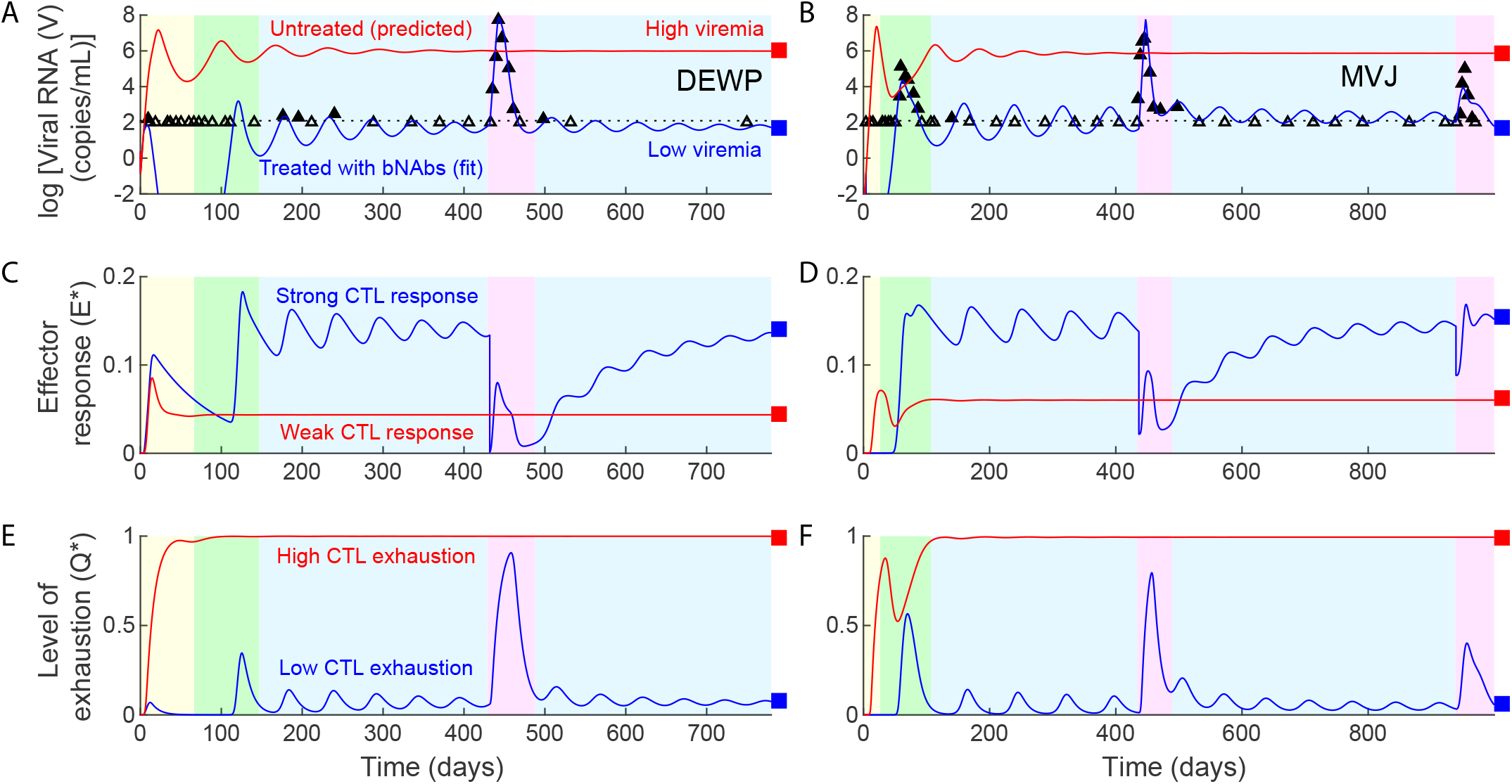
Early bNAb therapy induces a switch to viremic control. (**a,b**) Model fits (blue lines) to viral load data 10 (symbols) from the responder macaques DEWP and MVJ. (The parameter values used are in Tables 1 and S1). The corresponding dynamics of (**c,d**) the effector response and (**e,f**) the level of effector exhaustion. The phases are color coded as in Figure 2. Red lines in all panels indicate model predictions with the same parameter values but in the absence of bNAb therapy. Our model predicts thus that bNAb therapy switches disease dynamics from reaching the high viremic, disease progressive state to the state of viremic control. Black dashed lines in (**a,b**) indicate the viral load detection limit (100 RNA copies/mL).

Our model also predicts that early bNAb therapy drove the infection to the state of lasting viremic control (blue lines in Figure 6), in agreement with the observations in Nishimura *et al.*^10^. Further, our model predicts that pleiotropic effects of bNAbs were involved in orchestrating this transition: During acute infection, bNAbs suppressed the viremic peak by enhancing viral clearance. The lower viremia would limit effector exhaustion. Further, bNAbs would increase antigen presentation and effector stimulation. When the level of the administered bNAbs in circulation is diminished, viremia would resurge but in the presence of a primed effector pool, which would be larger and/or less severely exhausted than in untreated macaques (Figure 6c-f). The primed effector population was able to control the infection in our predictions. Further, based on our predictions, when the effector response becomes significant before bNAb clearance, viremia would rise minimally post bNAb therapy, as observed with the macaque DEWP (Figure 6, left). When the effector response is weaker, viremia would resurge post bNAb therapy, but the limited effector exhaustion together with increased effector stimulation from the residual bNAbs would trigger a strong enough effector response to reassert control, akin to the observations with the macaque MVJ (Figure 6, right).

The rise in viremia post treatment resembles an infection under conditions where the effector population is primed and thus has a head start over the virus. Indeed, our model calculations based on the hypothetical scenario where a higher effector population existed at the time of viral challenge showed spontaneous control (Figure 7).

**Fig. 7.**
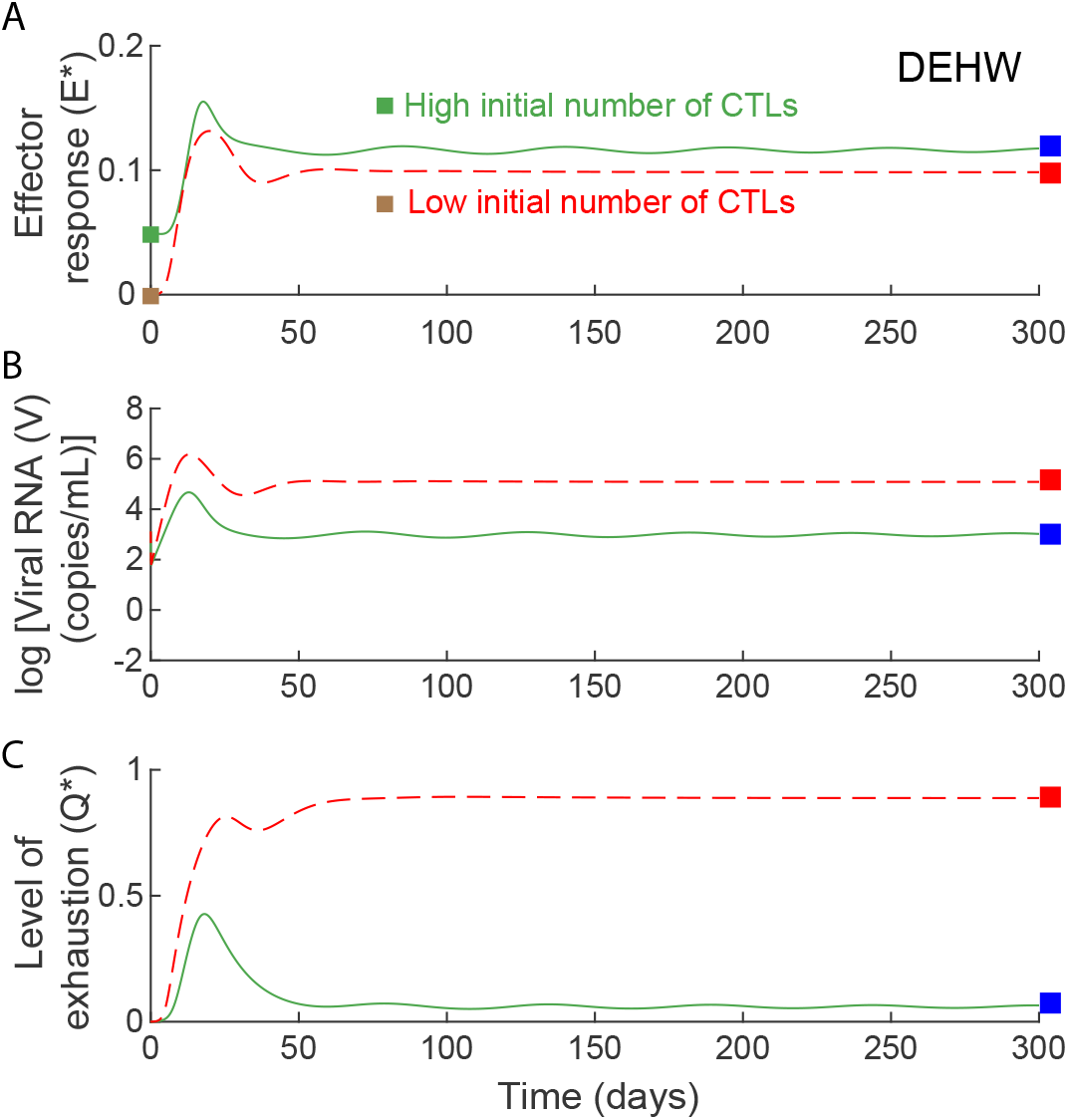
Model predictions of untreated infection dynamics with higher initial effector pool. Model predictions of effector, viral, and exhaustion dynamics for untreated infection show control with high (green) and progression with low (red) initial effector numbers (*E*\*(0)). All other parameters are the same as those that yield the best-fit to the macaque DEHW (Table S1).

Similarly, our model predicts that suppression of effectors due to anti-CD8 Abs up to a threshold after the establishment of control allowed resurgence in viremia but restored control, whereas suppression beyond the threshold sacrificed control and drove infection to the former, high viremic fate (Figure 2), explaining the difference between the controller macaques who reasserted viremic control following the transient rise in viremia upon the administration of anti-CD8 Abs and the one controller macaque (DFIK) that failed to reassert control and transitioned to the high viremic state ^10^. Note that the threshold can vary across macaques and corresponds, in our model, to the unstable boundary separating the stable states of viremic control and progressive disease (Figure 5).

When we repeated our analysis without either enhanced clearance of virus or upregulation of antigen presentation by bNAbs, our model failed to capture the observed viral load data robustly (Figures S2 and S3, and Tables S3 and S4). Specifically, neglecting bNAb-induced enhancement of viral clearance did not fit the data well (Figure S2), whereas neglecting bNAb-induced upregulation of antigen presentation yielded fits with a higher value of the Akaike information criterion (AIC) (Figure S3).

Our analysis suggests, thus, that the rapid clearance of the virus, which lowered viremia and reversed effector exhaustion, together with increased antigen uptake, which led to enhanced effector stimulation, resulted in the lasting control of viremia elicited by early, short-term bNAb therapy.

### bNAbs have an advantage over ART

If ART were used instead of bNAbs, for a duration equivalent to the time over which the administered bNAbs were in circulation, viral load quickly became undetectable during treatment in our model but rebounded to high levels post treatment (Figure 8), consistent with the 3 macaques administered ART in Nishimura *et al.*^10^. While both bNAb therapy and ART limited viremia (Figures 6a, 6b, 8a) and would thus prevent effector exhaustion (Figures 6e, 6f, 8c), bNAb therapy can additionally increase antigen presentation and effector stimulation (Figures 6c, 6d, 8b). The resurgence in viremia post the cessation of ART would then be akin to *de novo* infection, which typically reaches the high viremic fate. Only in the rare individuals in whom the latent reservoir is sufficiently small is this rise in viremia small enough for the effector population to catch up and establish control, explaining the critical role of the size of the latent reservoir in post treatment control with ART ^18^. The primed effector population following bNAb therapy, in contrast, drove the system to viremic control in our model. Because of their pleiotropic effects, bNAbs would thus have a significant advantage over ART, explaining their much higher success rate in achieving lasting control.

**Fig. 8.**
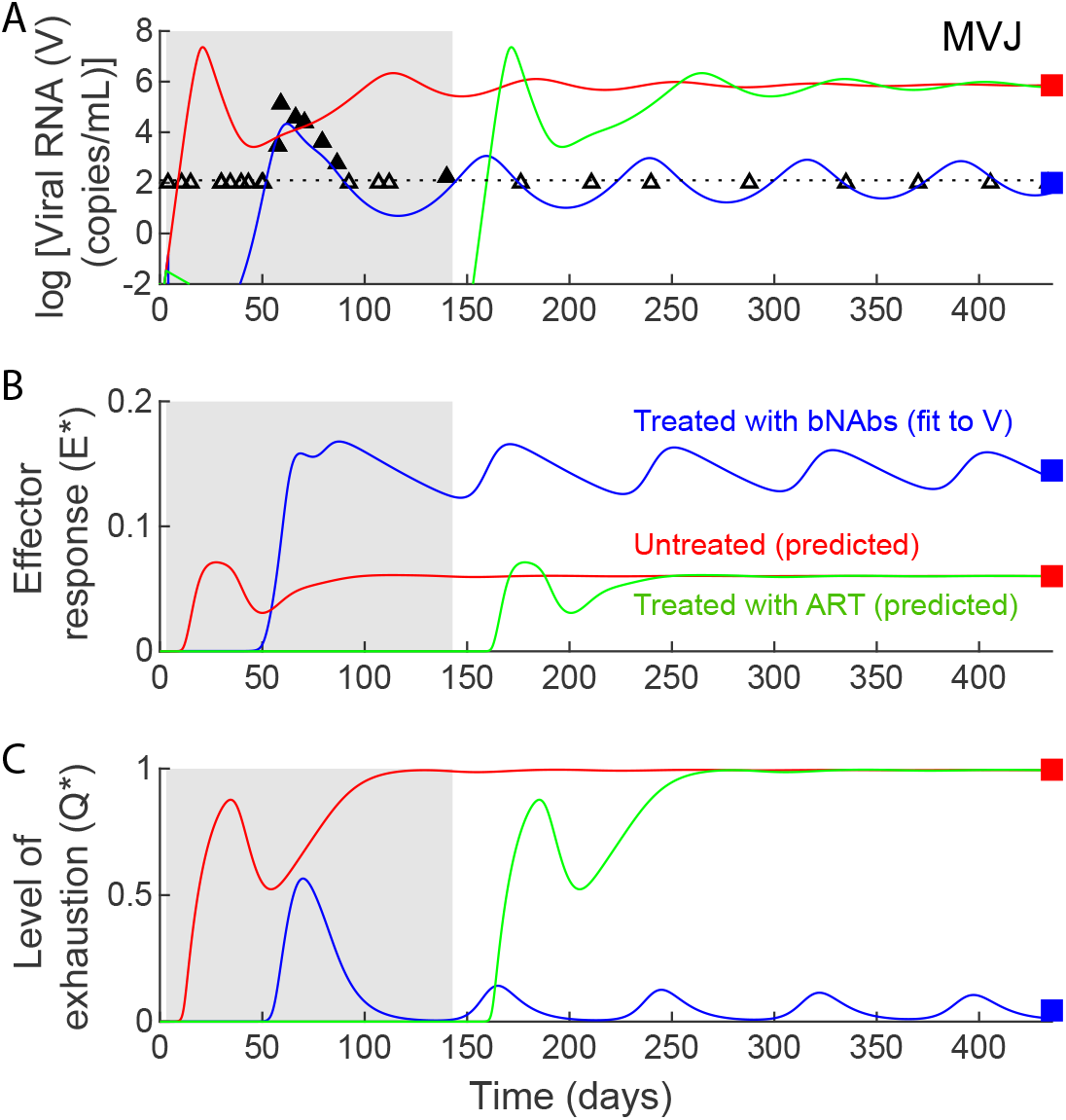
bNAbs succeed whereas ART fails to establish long-term viremic control. Simulated dynamics of (**a**) viral load, (**b**) effector response, and (**c**) effector exhaustion with ART (green lines) are shown in comparison with the corresponding dynamics without treatment (red lines) and with bNAb treatment (blue lines) using parameters that capture data for the macaque MVJ (symbols). (Parameter values used are in Tables 1 and S1. Efficacy of ART is assumed to be *ε* = 0.8.) The duration of ART is shown as a grey region. The black dashed lines in (**a**) indicates the viral load detection limit (100 RNA copies/mL).

## Discussion

The recent success of passive immunization with bNAbs in eliciting functional cure of HIV-1 infection can potentially revolutionize HIV-1 care. ^10–17^ In this study, using mathematical modelling and analysis of *in vivo* data, we present a dynamical systems view of the infection that offers an explanation of how early exposure to bNAbs induces lasting viremic control and does so better than ART.

Our analysis suggests that progressive disease and viremic control are states ‘intrinsic’ to the system. Interventions, with ART or bNAbs, can serve to alter the propensity with which the states are realized. In untreated infection, rapid viral growth can induce significant effector exhaustion and lead typically to progressive disease. ART suppresses viremia and would therefore at least in part reverse exhaustion. (Recent studies have argued that CTL exhaustion may not be fully reversible. ^39^) However, because successful ART can completely block viral replication, the absence of antigenic stimulation could eventually cause the activated effector population to fade. When viremia rises post treatment, if the activated effector population has faded significantly, the dynamics would mimic *de novo* infection leading to progressive disease. The resurgence of viremia post ART is due to the reactivation of latently infected cells. When the latent pool is small, this reactivation may lead to low viremia, which can stimulate effectors and culminate in viremic control. Indeed, a model developed to explain postART control found a critical dependence on the size of the latent reservoir.^18^ In contrast, bNAbs can additionally stimulate effectors by enhancing antigen presentation, a phenomenon first identified with cancers and termed the ‘vaccinal effect’ ^40^. Our model was most consistent with data when this effect was incorporated. When viremia rises post bNAb therapy, it then would encounter a primed effector population, which can drive the system to viremic control. Unless intense bNAb therapy completely halts active viral replication, akin to ART, the establishment of control is less likely to be sensitive to the latent pool size. Indeed, our model fit the macaque data without considering latently infected cells. bNAbs would therefore achieve lasting control in a far greater percentage of the population treated than ART.

Previous studies either examined the steady states alone^18^ or short-term viral load changes following exposure to bNAbs ^24^. Going beyond, our study captures the entire time course of viral load changes leading to lasting control, resulting in a more comprehensive understanding of functional cure. Our model predicts that through pleiotropic effects, bNAbs tilt the balance of the competing interactions between the virus and effectors in favour of effectors. It follows that other strategies that similarly tilt the balance may also elicit functional cure. For instance, immune checkpoint inhibitors can prevent effector exhaustion during infection. ^41^ Alternatively, effector cells may be adoptively transferred, a strategy that showed promise in a recent macaque study ^42^. Effector cells can also be stimulated using vaccines. ^43,44^ Our model predicts that a primed effector population at the time of viral challenge may prevent the infection from reaching the high viremic, progressive state and drive it instead to lasting viremic control, indicating a potentially favourable outcome of preventive T cell vaccines. Stimulating effectors using vaccines has been shown to suppress viral replication and lower the set-point viremia ^44,45^, but several additional design challenges must be overcome, including the need to stimulate effectors early enough and, interestingly, prevent effector exhaustion due to the vaccine^46^, for the successful deployment of T cell vaccines^47^. Further, such strategies may not work in children, where bNAbs may elicit control without effector responses, possibly due to the inadequately developed immune system. ^48^

Following the observation of Nishimura *et al.*^10^ that there was no significant generation of anti-gp120 Abs in controllers, we ignored the ability of bNAbs to upregulate the humoral response^49,50^. Our study is thus conservative in its assessment of the influence of bNAbs. We did not consider data from the three non-responder macaques, where functional cure was not established and CD4^+^ T cell counts gradually declined^10^. Describing the entire course of HIV infection from the acute phase to AIDS using a single model has been a long-standing challenge. ^51^ One possibility for the failure of bNAb therapy in the non-responders is that the bNAb therapy induced rapid viral load decline, limiting effector stimulation, and thus behaved like ART in the non-responders. ^10^ Alternatively, the non-responders may have had parameter settings that did not admit bistability, a possibility also with those who failed to achieve post-ART control^18^. Note that the determinants of post-ART control are yet to be established. ^8,9^ Finally, we did not consider viral evolution and resistance to bNAbs, recognizing that most controllers in Nishimura *et al.*^10^ saw no resistance. Future studies with other combinations of bNAbs may determine whether resistance must be accounted for in defining optimal bNAb treatments, especially for use during chronic infection where viral diversity is likely to be large ^52^.

In summary, our study presents the first quantitative description of viral dynamics following passive immunization with HIV-1 bNAbs; captures *in vivo* data; elucidates the mechanisms with which early, short-term bNAb therapy establishes lasting viremic control of SHIV infection; explains the advantages of bNAbs over ART; and suggests alternative avenues to induce functional cure of HIV-1 infection.

## Methods

### Model equations

We constructed the following equations to describe the viral dynamics depicted schematically in Figure 1.

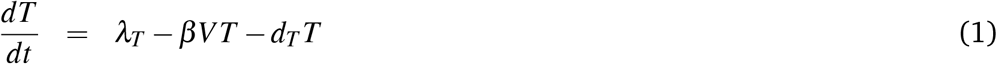

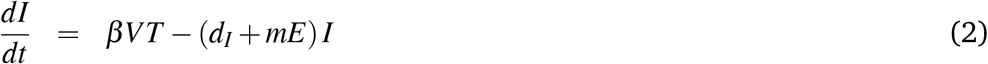

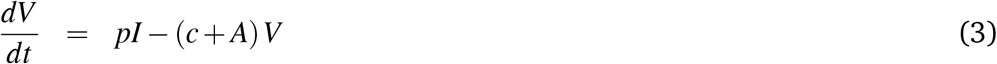

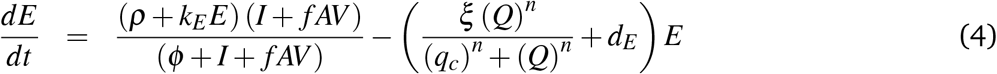

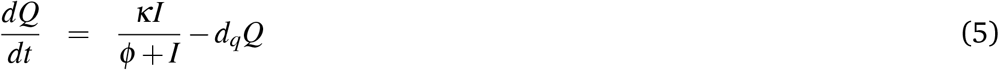

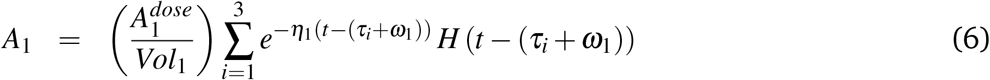

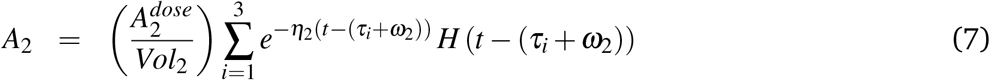

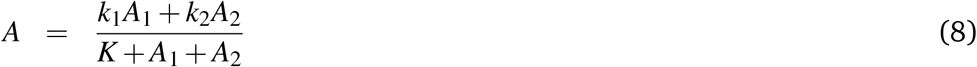

Here, uninfected target CD4^+^ T cells, *T*, are produced at the rate *λ*_*T*_, die at the per capita rate *d*_*T*_, and are infected by virions, *V*, with the second order rate constant *β*, yielding productively-infected cells, *I*. Cells *I* die at the per capita rate *d*_*I*_due to viral cytopathicity and are killed by effector cells, *E*, with the second order rate constant *m*. Cells *I* produce free virions at the rate *p* per cell. The virions are cleared in the absence of administered bNAbs with the rate constant *c*. Administered bNAbs are assumed to induce a net enhancement of the viral clearance rate by a time-dependent amount, *A*, which is a saturable function of the instantaneous serum concentration of the two bNAbs, 3BNC117 (*A*_1_) and 10-1074 (*A*_2_). The functional form mimics the 2D Hill equation derived recently to describe the effect of drug combinations^53,54^ and has been used to quantify combinations of Abs and virus entry inhibitors ^54,55^, expected to be similar to the combination of bNAbs targeting non-overlapping sites on the HIV-1 envelope used here ^10^. The net enhancement of viral clearance combines the direct effect on viral clearance by bNAbs ^22^ as well as the reduction in infectivity, *β*, due to viral neutralization by the bNAbs ^56^. By preventing new infections, virus neutralization lowers the number of infected cells and hence overall viral production. Because viral production and clearance are rapid compared to other processes^57^, the effect of lower viral production on viral dynamics is indistinguishable through viral load measurements from the effect of enhanced viral clearance. Indeed, the pseudo-steady state approximation applied to Eq. (3) yields *V* ≊ *pI*/(*c* + *A*), which when substituted into Eqs. (1) and (2) amounts to a reduction in *β* by the factor (*c* + *A*)*/c*. Conversely, a reduction in *β* can be subsumed into an effect on *c*.

Administered bNAbs also bind to virions and increase antigen uptake by antigen presenting cells (APCs), which in turn would increase stimulation of effectors. If we define *P* as the population of activated APCs, which can thus stimulate effectors, then we may write *dP/dt* = *γ I* + *νAV* − *d_P_P*, where *γ* is the rate constant of APC stimulation by infected cells, *d*_*P*_ their per capita death rate, and *σ* the fractional rate of opsonized virions cleared (AV) that is taken up by APCs. We assume naïve APCs to be in large excess. Assuming pseudo steady state yields *P* = (*γ I* + *νAV*) */d_P_*. Finally, letting effector cell stimulation be a saturable function of *P* and simplifying yields the expression in Eq. (4) for effector stimulation by antigen. In effect, the total serum antigen available for presentation to effectors, *I* + *f AV*, stimulates immune cells such as naïve CD8^+^ T cells ^19,58^, NK cells ^59^, and others, assumed not to be limiting, into antiviral effectors, *E*, with the maximal rate, *ρ*, and the half-maximal antigen-sensitivity parameter, *ϕ*_1_. (Here, *f* = *ν/γ*.) We note that *E* is a composite effector pool consisting of SHIV-specific CTLs, NK cells, and other effectors. ^10,18,60^ The saturation of the stimulation with increasing levels of total serum antigen is reflective of limitations in cellular interactions processes, such as the time to find interacting partners, the duration of each interaction, and cytokine/chemokine signalling. ^59,61–63^ Saturating forms are argued to be more realistic than mass-action forms^62^ and follow from kinetic considerations of these interactions under the total quasi-steady state approximation^63^. Following activation, effectors such as CTLs and NK cells enter a proliferation program that does not require antigen but can be modulated by antigen, cytokines and other stimulatory and co-stimulatory signals. ^59,64–67^ Accordingly, we let effectors proliferate ^68^ with the rate constant *k*_*E*_, and, for simplicity, subject to the same saturating dependence on antigen as activation. Effectors suffer exhaustion ^19,69^ at the maximal rate *ξ*, with the half-maximal parameter *q*_*c*_and Hill coefficient *n*. The level of exhaustion, *Q*, increases with antigen level at the maximal rate *κ* and with the half-maximal parameter *ϕ*_2_. Exhaustion is reversed with the rate constant *d_q_*. effectors are lost at the per capita rate *d*_*E*_. We note that *E* is a measure of the effective effector pool, consisting of activated NK cells and SHIV-specific CTLs, and factoring in the overall level of exhaustion, *Q*. *E* is thus not to be viewed as a cell count. Similarly, *Q* is not a measure of the pool of exhausted cells. Our model of effector stimulation and exhaustion mimics an earlier study.^19^ An alternative model, developed to describe post-ART control of HIV ^18^, also shows bistability but could not fit the present data with bNAb therapy (Figure S4 and Table S5). Similarly, variants of our model without the Hill coefficient, i.e., n = 1 ^18^ (Figure S5 and Table S6), and with a Hill coefficient of n = 3 ^19^ (Figure S6 and Table S7), or without an explicit effector response (Figure S7 and Table S8) could not fit the data. We also fit the viral load data by letting *k*_*E*_ vary. The model fits (Figure S8 and Table S9) were comparable to the main model (Figure 2) but with higher AIC (Table 3).

**Table 3.**
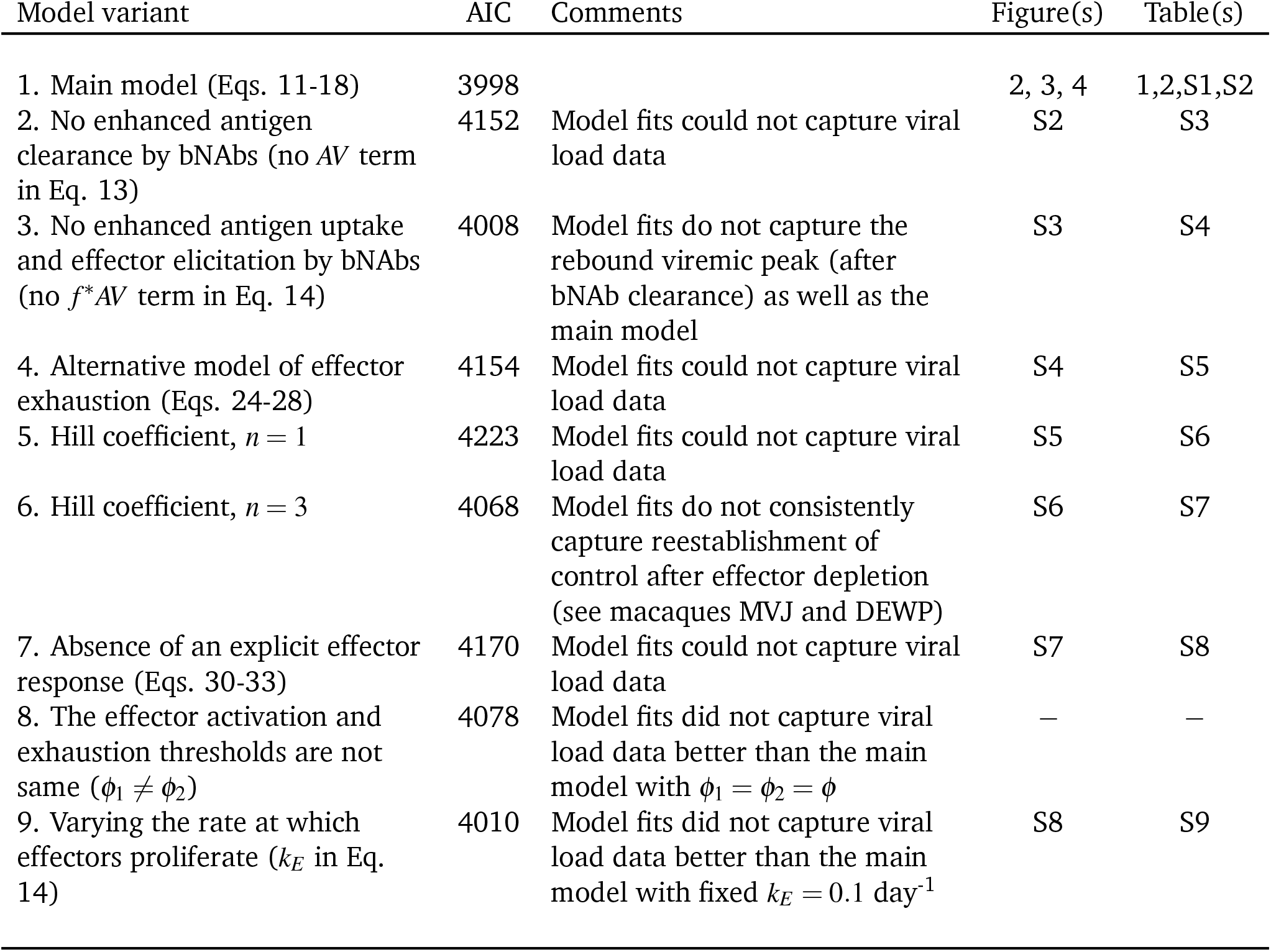
Summary table comparing the Akaike information criterion (AIC) of the main model with model variants.

To mimic the dosing protocol in Nishimura *et al.*^10^, we let the serum concentration levels of the bNAbs, *A*_1_ and *A*_2_, rise by the extent 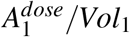 and 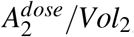, immediately upon the administration of a bNAb dose, and decline exponentially with the rate constants *η*_1_ and *η*_2_, respectively. Here, 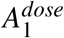and 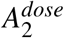 correspond to the dosages of the two bnAbs, and *Vol*_1_ and *Vol*_2_ correspond to the bNAb-specific volumes of distribution. The three doses were administered on days *τ*_1_ = 3, *τ*_2_ = 7, and *τ*_3_ = 10, respectively. Additionally, we allowed a delay in bNAb action, *ω*_1_ and *ω*_2_, following each dose, to account for any pharmacological effects within individual macaques. The Heaviside function, *H*(*t* < (*τ*_*i*_+ *ω_j_*)) = 0 and *H*(*t* ≥ (*τ*_*i*_+ *ω_j_*)) = 1, accounts for bNAb dynamics based on these dosing times. The maximal efficacies of the respective bNAbs, achieved when they are in excess, are set to *k*_1_ and *k*_2_, respectively, and their half-maximal efficacy is defined the Hill function threshold *K*. To compare the influence of bNAbs with that of ART, we assumed that ART reduces *β* by a factor 1 − *ε*, where *ε* is the efficacy of the drug combination^18^.

Long after bNAbs were cleared from circulation, Nishimura *et al.*^10^ administered anti-CD8*α* antibodies to some macaques, which resulted in the depletion of effectors such as CD8^+^ T, NK, NKT, and *γδ* T cells. Nishimura *et al.*^10^ also administered anti-CD8*β* antibodies to some macaques, which presumably only depleted CD8^+^ T cells. To describe the resulting changes in viremia, we assumed that the depleting effects of anti-CD8*α* and anti-CD8*β* antibodies started at time points *θ*_*α*_ and *θβ*, respectively, at which points we reduced the effector populations by fractions *ζ*_*α*_ and *ζ_β_*. Further, we assumed that anti-CD8*α* antibodies neutralized all host effector functions, which we modelled by setting *m* = 0 for a duration *θ*_*m*_ representing the residence time of the depleting antibodies.

The procedures for solving the above model equations, parameter estimation, and data fitting are described next. All the data employed for fitting was obtained by digitizing the data published in Nishimura *et al.*^10^ using the software Engauge digitizer, and is available as a supplementary excel file (Dataset.xlsx).

### Solution of model equations

For ease of solution and parameter estimation, we rescaled Eqs. 1–8 in the main text using the quantities in Eqs. 9 and 10 and obtained an equivalent model with fewer parameters (Eqs. 11–18). bNAb concentrations, *A*_1_ and *A*_2_, and viral load, *V*, were not scaled because they were used for fitting data.

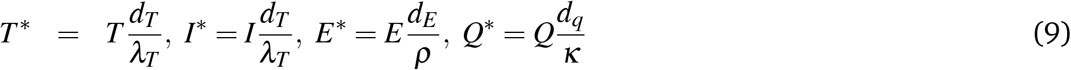

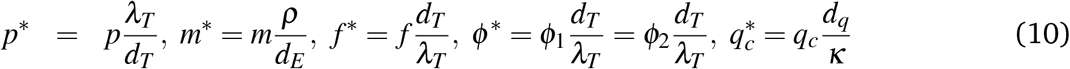

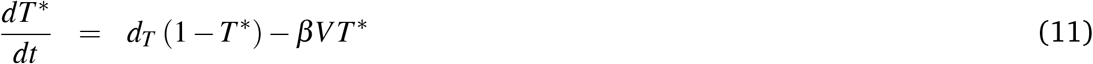

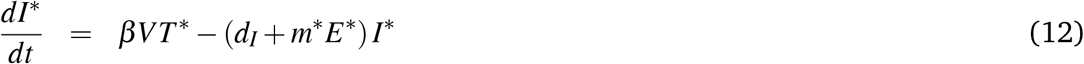

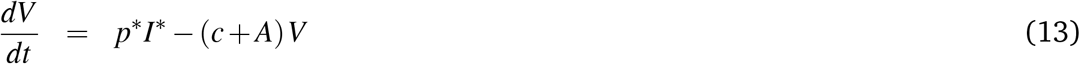

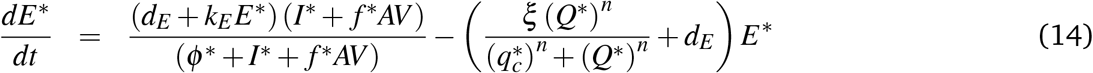

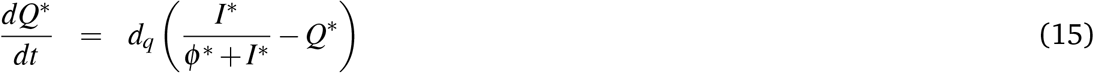

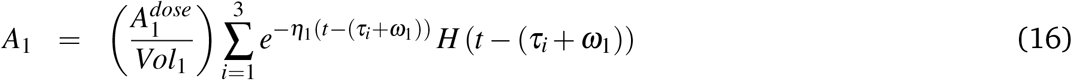

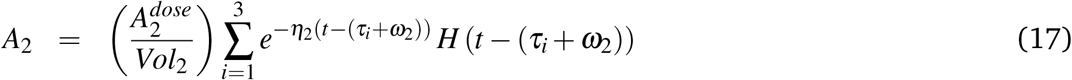

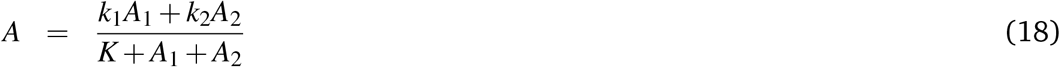

The model equations were integrated using the initial conditions *T* *(0) = 1, *I*\*(0) = *E*\*(0) = *Q*\*(0) = 0,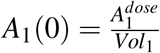 and 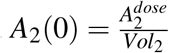. The initial viral load, *V* (0), varied for individual macaques and was estimated from fits.

We also obtained steady state solutions by setting the left-hand sides of all equations to zero and solving the resulting coupled non-linear equations in Mathematica. We subsequently performed linear stability analysis, by computing the eigenvalues of the Jacobian matrix of the above dynamical system, to assess the stability of the steady states.

### Parameter estimates and data fitting

The data available from Nishimura *et al.*^10^ for fitting this model consists of longitudinal values of the plasma SHIV viral load (*V*) and serum bNAb concentrations (*A*_1_ and *A*_2_). For this model, we performed formal identifiability analysis of the model parameters by employing the Exact Arithmetic Rank (EAR) implementation in Mathematica (IdentifiabilityAnalysis package^70^), assuming that measurements of *V*, *A*_1_ and *A*_2_ are perfect and continuous in time. The analysis revealed that upon fixing 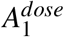 and 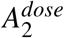, whose values can be estimated from Nishimura *et al.*^10^, all the other parameters in the model are fully identifiable from the data.

Wherever possible, we fixed model parameter values based on previously published estimates (Table 1). For instance, following Conway and Perelson ^18^, we fixed the death rate of target cells, *d*_*T*_ = 0.01 day^−1^. We let the death rate of infected cells due to viral cytopathicity be *d*_*I*_= 0.5 day^−1^, which is ~ 50% of the mean overall death rate of infected cells ^61^, and lies at the lower end of the range of the overall death rate, 0.5 − 1.7 day^−1^ for HIV-1^71^, with the rest attributed to effector killing. The viral clearance rate was estimated ^18,72^ to be *c* = 23 day^−1^ for HIV-1, but has been shown to be much higher (*c* = 38 − 302 day^−1^ for SIV ^22,29,30^. We conservatively chose a value of *c* = 38 day^−1^. To estimate the effector expansion rate, *k*_*E*_, we obtained fits w ith *k*_*E*_ adjustable, which yielded best-fit *k*_*E*_ ~ 0.3 day^−1^. The model, however, had a higher value of the Akaike Information Criterion (AIC) than when *k*_*E*_ was fixed (Table 3). Here, we therefore chose a model with fixed *k*_*E*_ and conservatively set *k*_*E*_ = 0.1 day^−1^. (Note that the *k*_*E*_ here represents the effective expansion of all effectors and not CTLs alone, which may have a higher expansion rate. ^18^) We set the Hill coefficient for exhaustion, *n* = 4, following studies that recognize underlying non-linearities ^18,19^ and because smaller integral values of *n* did not yield good fits (see below). Further, we set *ϕ*_1_ = *ϕ*_2_ = *ϕ* following earlier studies ^19^ and because fitting them as separate variables yielded best-fit values that were close, *ϕ*_1_ ~ 2.1 × 10^−5^ and *ϕ*_2_ ~ 5.7 × 10^−5^, but with a higher AIC (Table 3).

We divided the remaining parameters into two sets, *ϑ* = {*β*, *m*\*, *p*\*, *f* *, *ϕ* *, *d*_*E*_, *ξ*, *V* (0), *Vol*_1_, *Vol*_2_, *k*_1_, *k*_2_, *K*} and *ρ* = {*ω*_1_, *ω*_2_, *η*_1_, *η*_2_}, the former assumed to follow log-normal and the latter logitnormal distributions, respectively. To estimate these parameters, we employed a population-based fitting approach using non-linear mixed effects (NLME) models to jointly fit the log plasma viral load (both treated and untreated animals) and antibody concentrations of the ten responder macaques in Nishimura *et al.*^10^ which do not exhibit consistent CD4^+^ decline: DEMR, MVJ, DEWP, DEWL, MAF, DFIK, DFKX, DFFX, DEHW, and DEBA. (A supplementary excel file, *Database.xlsx*, containing digitized viral loads and serum concentrations of bNAbs 3BNC117 and 10-1074 with time for all bNAb-treated and untreated macaques from Nishimura *et al*^10^ is available with this manuscript.) Briefly, the parameters for all the individual macaques were assumed to be sampled from a common population distribution, and the aim of the fitting exercise was to obtain estimates for the mean and variance of this distribution for each parameter. For each macaque *i*, parameters *ϑ*_*i*_were estimated assuming underlying log-normal distributions of the form:

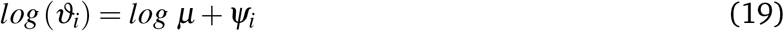

where *μ* is the set of the population means and *Ψ*_*i*_~ *N*(0*, σ*) are normally distributed ‘random effects’ whose variances are to be estimated. Similarly, parameters *ρ*_*i*_ for macaque *i* were estimated assuming underlying logit-normal distributions of the form:

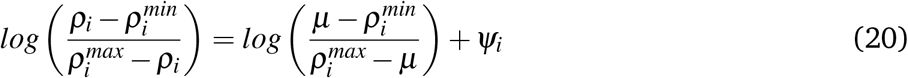

where 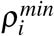 and 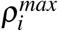 are the minimum and maximum allowed values for parameters in *ρ_i_*. The latter limits were set so that *ω*_1_*, ω*_2_ ∈ [0, 5], and *η*_1_*, η*_2_ ∈ [0.01 − 7] day^−1^, expected from the 1 week dosing interval employed. To account for measurement errors in observed viral loads and bNAb levels (*y*(*t*)), we defined a ‘combined error’ model such that *y*(*t*) is normally distributed around a true value (*y*\*(*t*)) as described by the following equation:

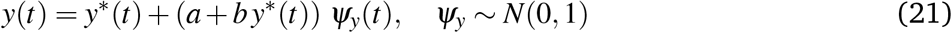

where *a* is the constant error term and *b* is the error term proportional to the true value, *y*\*(*t*).

All fitting except CD8 depletion was performed using Monolix software version 2019R1 (www.lixoft.eu), where the population estimates were obtained by maximum likelihood estimation. Untreated monkeys infected with the SHIV_AD8-EO_ viral strain consistently proceed towards high viremic chronic infection and immunodeficiency^37,73,74^. To recapitulate this behaviour, we simultaneously solved the model without bNAbs and ensured that the set point viral load attained values greater than 10^3.5^ copies/mL (enforced using the ‘censoring’ keyword during data input in Monolix). Parameter exploration was carried out using the stochastic approximation using expectation maximization (SAEM) algorithm implemented in Monolix^75^, and we simulated a large number of realizations (100 − 200) to produce consistent final parameter estimates that were biologically realistic. Comparisons of some of the population parameters with independent estimates from previous studies further validated our fitting procedure. For instance, the estimated population value of *p*\* corresponds to a burst size of 6595 virions per infected cell (Table 2), consistent with current estimates of ~ 10^4^ virions per cell^76^ given a target cell production rate ^18^ of *λ*_*T*_ = 10^4^ cells mL^−1^ day^−1^. Similarly, the estimated infectivity, *β* = 10^−8^ virion^−1^ mL day^−1^, is close to the previous estimate of 2.4 × 10^−8^ virion^−1^ mL day^−1 71^. Finally, we note that the enhanced clearance rate of virions, *c* + *A*, has a maximum of ~ 145 day^−1^, which is ~ 3.8-fold above the natural clearance rate, *c* = 38 day^−1^, consistent with the 3 − 4 fold increase in clearance rate due to bNAbs observed in other studies ^22^.

For capturing CD8^+^ T cell depletion experiments, since explicit dynamics of the anti-CD8 antibodies were not available from Nishimura *et al.*^10^, simultaneous fitting with Monolix consistently yielded poor fits to viral load data, especially in phase II (green region in Figure 2), often with unrealistic parameter estimates. Therefore, we first obtained parameters for each monkey from the population fits without CD8 depletion, and subsequently, using these parameters, we fit viral loads upon CD8^+^ T cell depletion to individual monkey data. Briefly, Nishimura *et al.*^10^ administered anti-CD8*α* antibodies to 6 macaques (MVJ, DEMR, DEWL, DEWP, MAF and DFIK) that resulted in depletion of CD8^+^ T, NK, NKT, and *γδ* T cells. All macaques exhibited a sharp increase in viremia, followed by reassertion of control in 5 macaques, and loss of control in one macaque (DFIK). Nishimura *et al.*^10^ also administered anti-CD8*β* antibodies to 3 macaques (MVJ, DEMR and DEWL) that only depleted CD8^+^ T cells, resulting in a smaller increase in viremia compared to the administration of anti-CD8*α* antibodies. We captured these observations through model predictions for macaques MVJ, DEMR, DEWL, and MAF. We assumed that the depleting effects of anti-CD8*α* and anti-CD8*β* antibodies started at time points *θ*_*α*_ and *θβ*, respectively, at which points we reduced the effector populations by fractions *ζ*_*α*_ and *ζβ*. Further, we assumed that anti-CD8*α* antibodies neutralized all host effector functions, which we modeled by setting *m* = 0 for a duration *θ*_*m*_ representing the residence time of the depleting antibodies. We fit *θ*_*α*_, *θβ*, *θ_m_*, *ζ*_*α*_, and *ζ_β_* to viral load data following the administration of depleting antibodies from all macaques (best-fits in phase IV of Figure 2). Fits were obtained using the ‘multistart’ and ‘lsqcurvefit’ tools along with the ode15s solver in Matlab (`in.mathworks.com`).

### Model variants

Many model variants were tested for their ability to accurately capture the data with fewer parameters.

#### Specific bNAb effects

We tested variants of our model without specific bNAb mediated effects such as enhanced antigen clearance (no *AV* term in Eq. 13) or without enhanced antigen uptake and subsequent effector elicitation (no *f* \**AV* term in Eq. 14). The model without antigen clearance failed to capture the viral load patterns (Figure S2) while the model without enhanced effector elicitation yielded poorer fits (Figure S3) but with higher AIC (Table 3).

#### Effector exhaustion

We also tested a simpler model of exhaustion, previously reported by Conway and Perelson^18^, where the loss of effector cells due to exhaustion is a saturating function of the antigen density. The scaled model equations are:

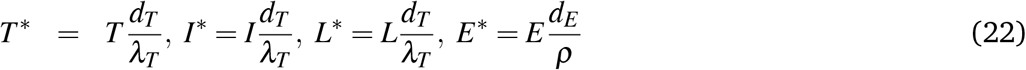

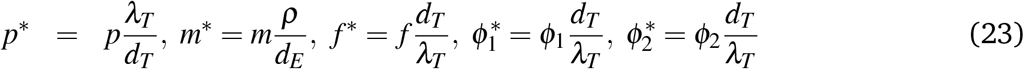

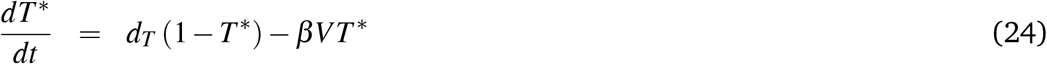

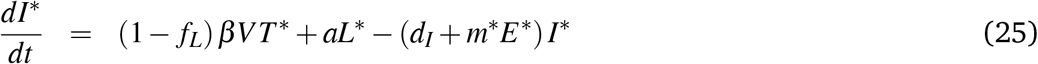

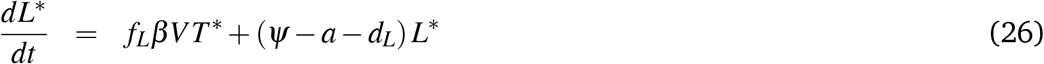

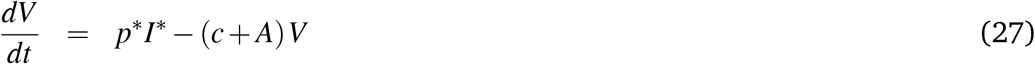

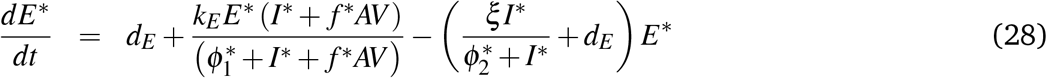

bNAb pharmacodynamics (*A*_1_ and *A*_2_) remained the same as Eqs. 16-18. *ϕ*_1_ and *ϕ*_2_ are the effector activation and exhaustion thresholds, respectively. In this model, a fraction (*f_L_*) of uninfected CD4^+^ infections yield latently-infected cells ^77^, *L*, and the rest, productively-infected cells, *I*. Cells *L* proliferate, get activated, and die at the per capita rates *Ψ*, *a*, and *d_L_*, respectively. We again followed the population-based fitting strategy above. The model failed to capture the viral load patterns observed within the 200 stochastic realizations employed (Figure S4).

#### Hill coefficient

The Hill coefficient for exhaustion, *n*, which dictates the sharpness of the exhaustion switch, cannot be accurately estimated from the data. While we use *n* = 4 (Eq. S6), previous studies have used values of 1 and 3^18,19^. Therefore, by employing the same population-based fitting procedure detailed above and with 100 − 200 realizations per model, we fit *V*, *A*_1_ and *A*_2_ to model variants with Hill coefficients *n* = 1 and *n* = 3. While the model with *n* = 1 failed to capture the viral loads (Figure S5), with *n* = 3, viral load patterns upon early bNAb therapy but not effector depletion (Figure S6) were captured. Specifically, upon effector depletion with anti-CD8*α* and anti-CD8*β* antibodies for monkeys DEWP and MVJ, respectively, while measurements by Nishimura *et al.,*^10^ showed reestablishment of control, fits exhibited loss of control and attainment of the high viremic steady state. *Absence of cytotoxic effector cells.* We checked whether a basic viral dynamics model, with the latent pool but without effectors could capture the macaque viral load data. The scaled model equations are:

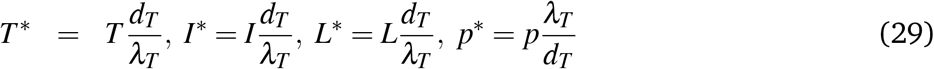

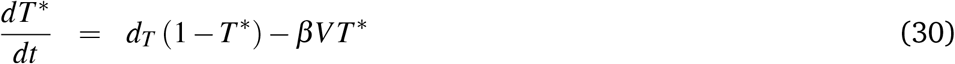

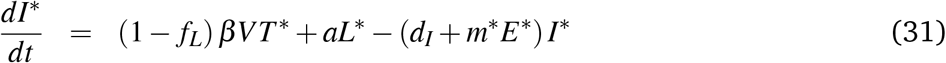

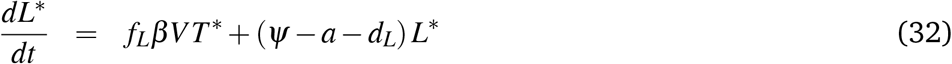

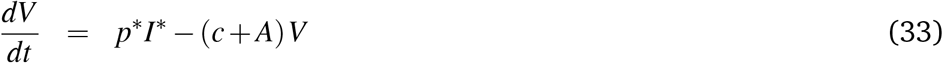

where bNAb pharmacodynamics (*A*_1_ and *A*_2_) remain as before (Eqs. S9 and S10). Again, the model could not capture the *in vivo* temporal viremic patterns (Figure S7), reaffirming the role of the effector response in establishing robust viremic control.

#### Effector proliferation rate

Lastly, we checked whether a varying effector proliferation rate (*k*_*E*_ in Eq. 14) made a difference to the model fits to viral load data. The fits (Figure S8) were not better than those obtained with our main model by fixing *k*_*E*_ = 0.1 day^−1^ (Figure 2), but had higher AIC (Table 3).

## Acknowledgements

We thank Alan Perelson, Rustom Antia, and Pranesh Padmanabhan for comments, and Fabian Cardozo-Ojeda for help with data analysis.

## Author contributions

Conceptualization - RD, RR and NMD; Methodology - RD and RR; Software - RR and RD; Validation - RD, RR and NMD; Formal Analysis - RD, RR and NMD; Investigation - RD, RR and NMD; Resources - RD, RR and NMD; Data Curation - RR and RD; Writing (original draft) - RD and NMD; Writing (review and editing) - RD, RR and NMD; Visualization - RR and RD; Supervision - RD and NMD; Project Administration - RD, RR and NMD; Funding Acquisition - NMD.

