## Supplementary Information for "Early exposure to broadly neutralizing antibodies may trigger a dynamical switch from progressive disease to lasting control of SHIV infection"

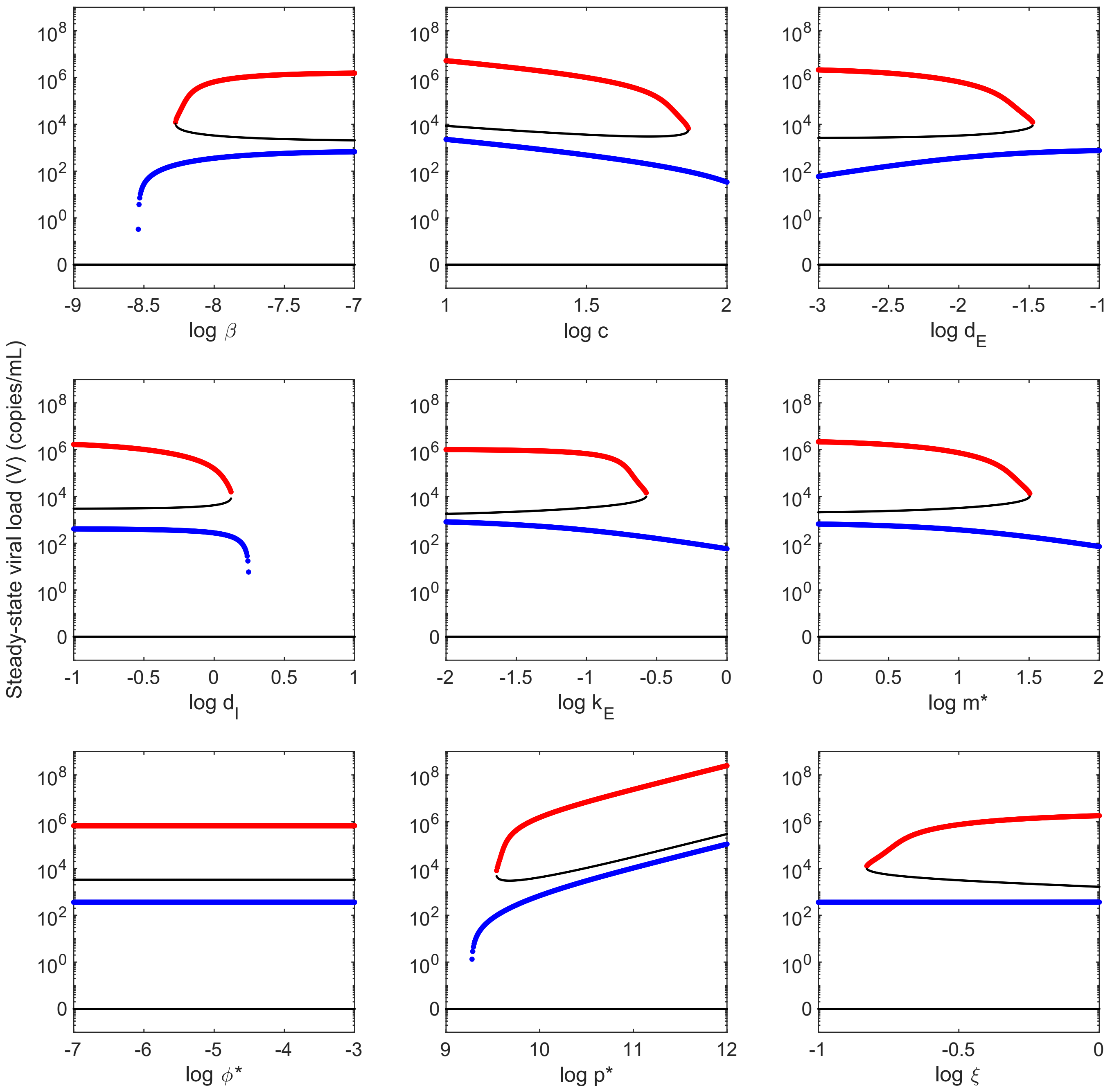


**Fig. S1 Bistability and bifurcation diagrams.** Steady states of our model (see Methods) obtained by varying underlying parameters (different panels) one at a time over wide ranges about their values listed in Tables 1 and 2. The stable states of high and low viremia are shown in red and blue, respectively. Black lines represent unstable steady states. Parameters: $\beta$ - infectivity of virions, $c$ - viral clearance rate, $d_E$ - death rate of effectors, $d_I$ - death rate of infected CD4$^+$ T cells, $k_E$ - proliferation rate of effectors, $m^*$ - rate at which effectors kill infected cells, $\phi^*$ - threshold for effector activation as well as level of exhaustion, $p^*$ - burst size, and $\xi$ - maximal rate of effector exhaustion.

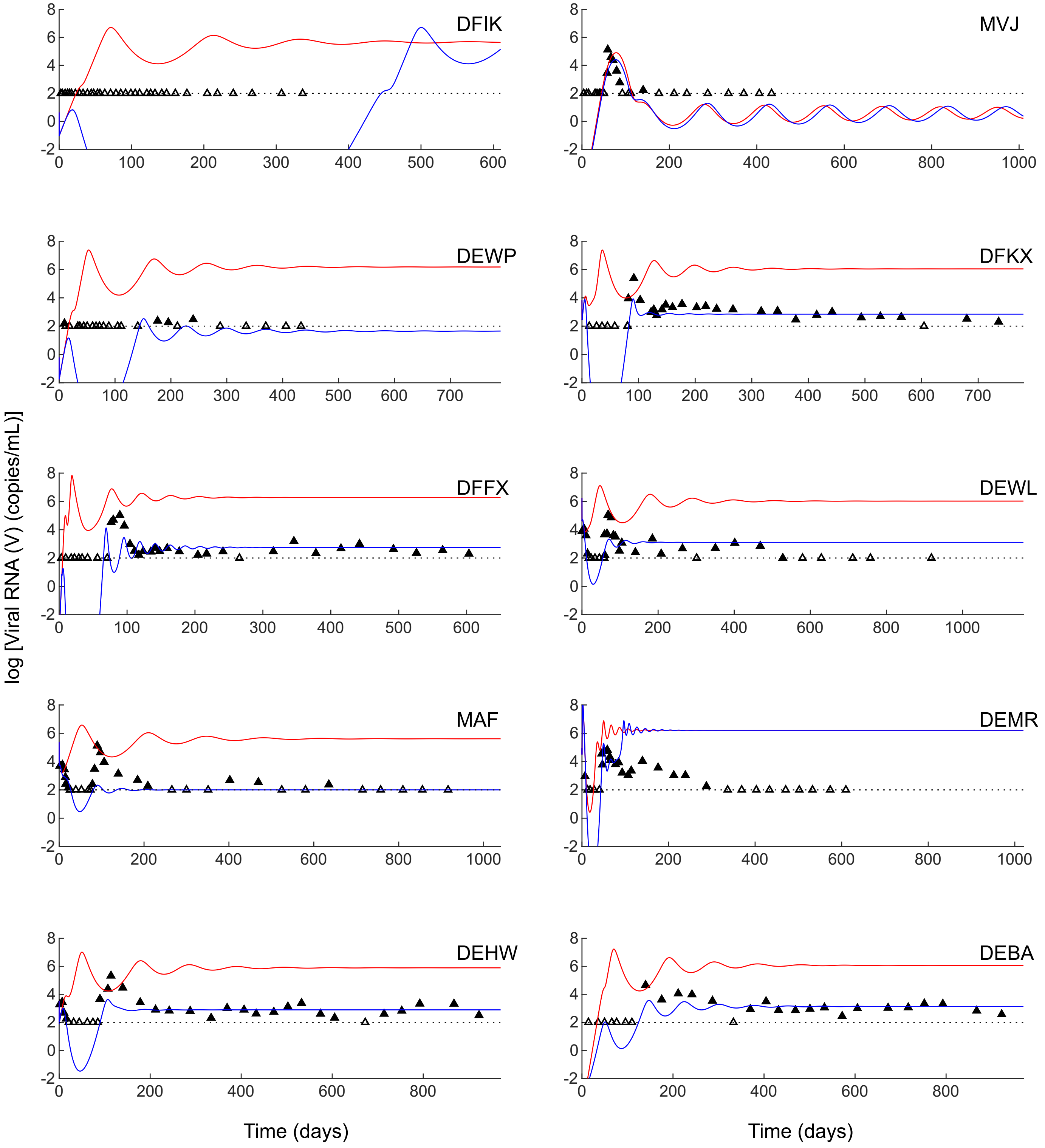


**Fig. S2 Fits of model without enhanced antigen clearance by bNAbs.** Fitting our model (Eqs. 11-18) without enhanced antigen clearance by bNAbs (no $AV$ term in Eq. 13) following the procedure outlined in the Methods yielded poor fits (blue) to the data (parameters in Table S3). Corresponding predictions without treatment are in red.

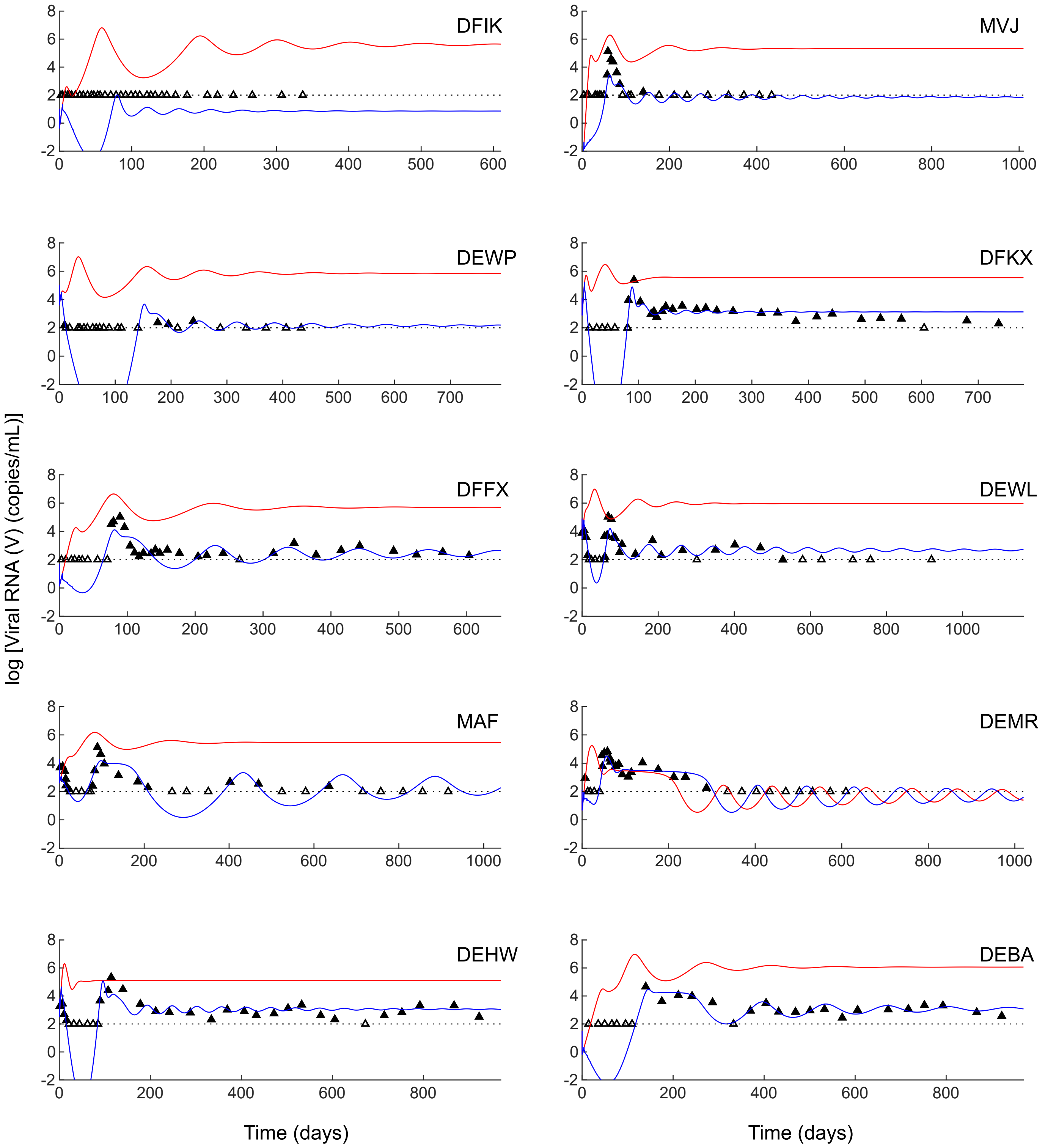


**Fig. S3 Fits of model without enhanced effector elicitation by bNAbs.** Fitting our model (Eqs. 11-18) without enhanced antigen uptake and subsequent effector elicitation by bNAbs (no $f^{*}AV$ term in Eq. 14) following the procedure outlined in the Methods yielded poorer fits (blue) to the data (parameters in Table S4) compared to the main model (Figure 2) and yielded a higher AIC (Table 3). Corresponding model predictions without treatment are in red.

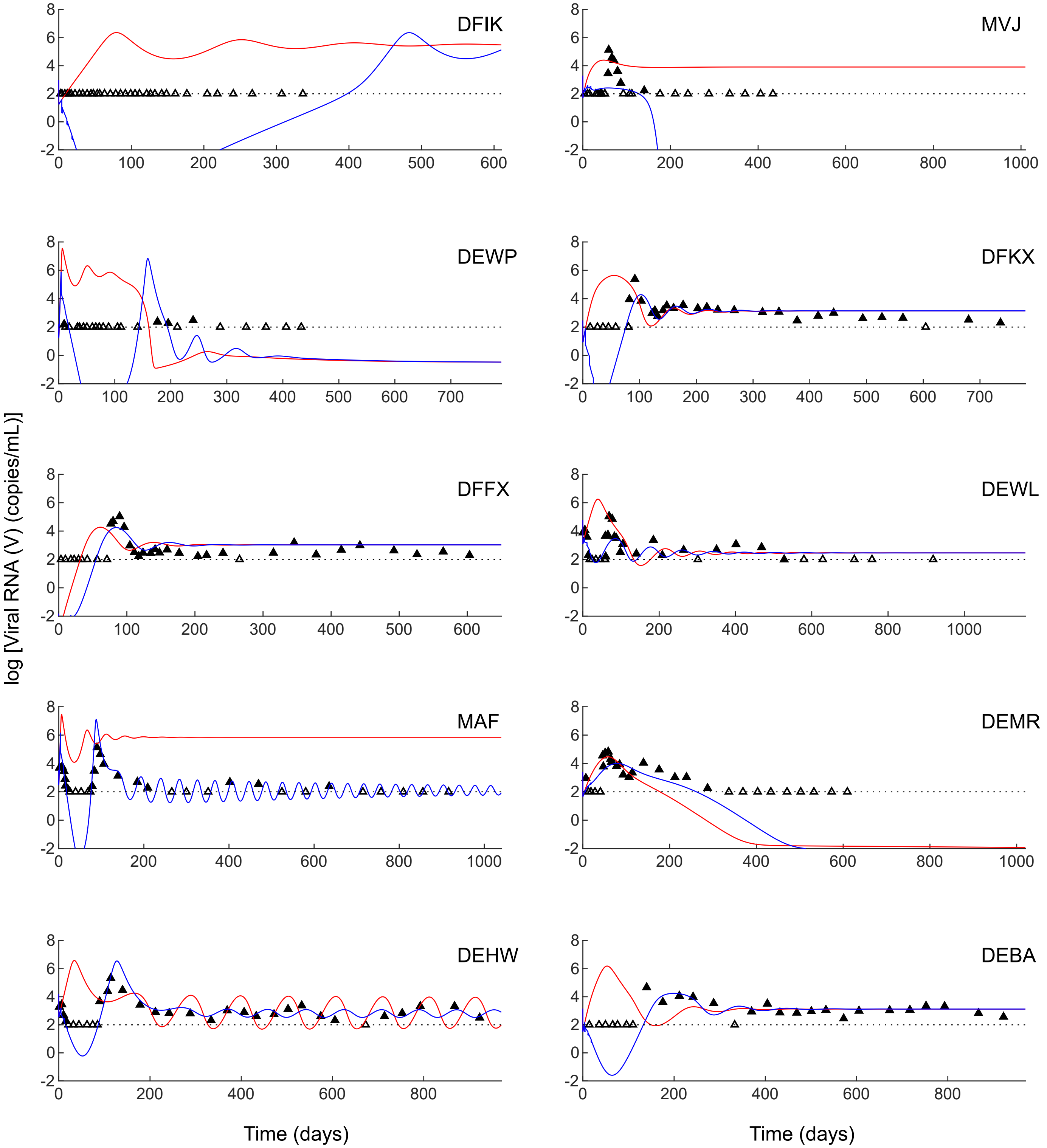


**Fig. S4 Alternative model of effector exhaustion.** Fitting with alternative model of effector exhaustion (Eqs. 24-28) following the procedure outlined in the Methods yielded poor fits (blue) to the data (parameters in Table S5). Corresponding predictions without treatment are in red.

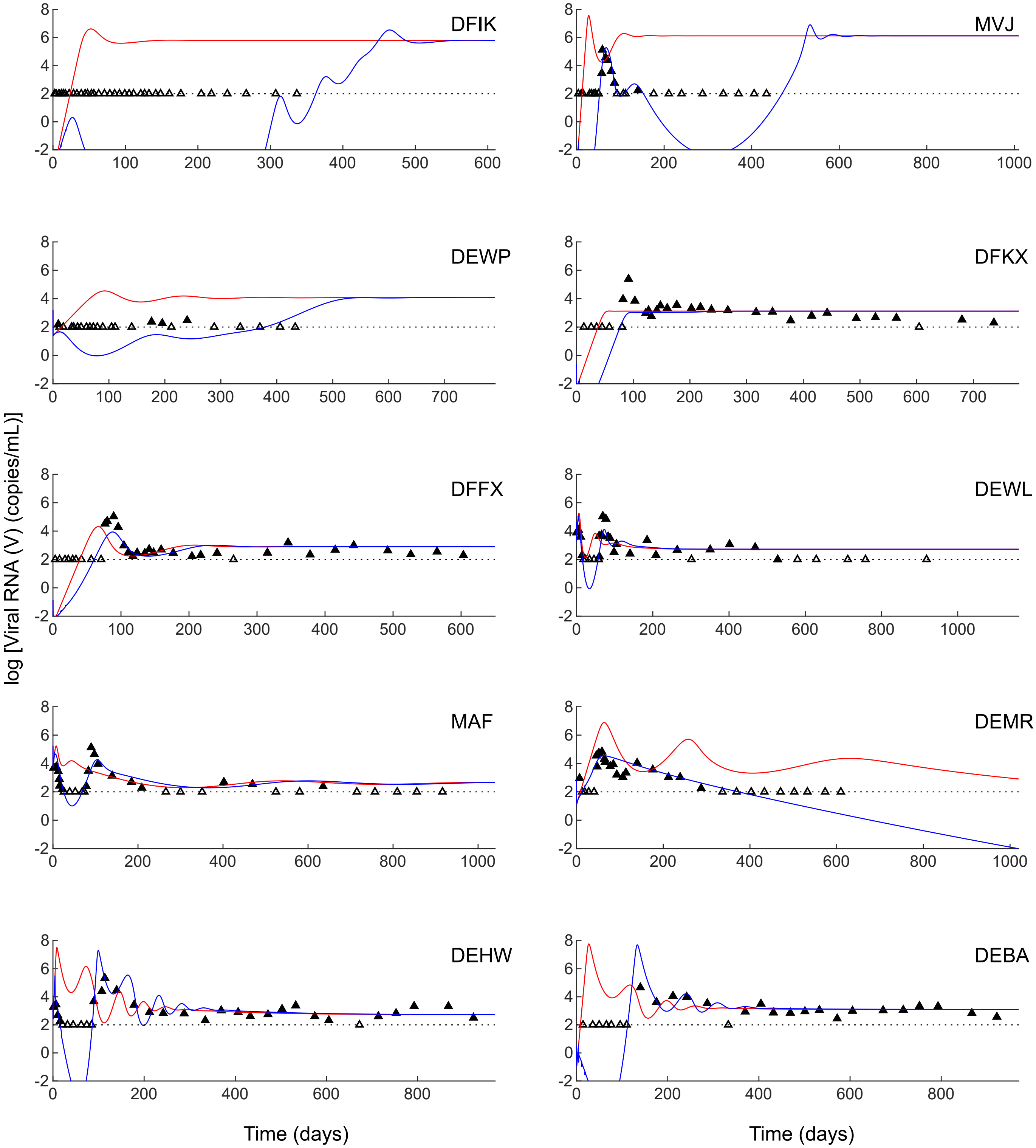


**Fig. S5 Hill coefficient,** $n = 1$. Fitting our model (Eqs. 11-18) with a Hill coefficient $n = 1$ following the procedure outlined in Methods yielded poor fits (blue) to the data (parameters in Table S6). Corresponding predictions without treatment are in red.

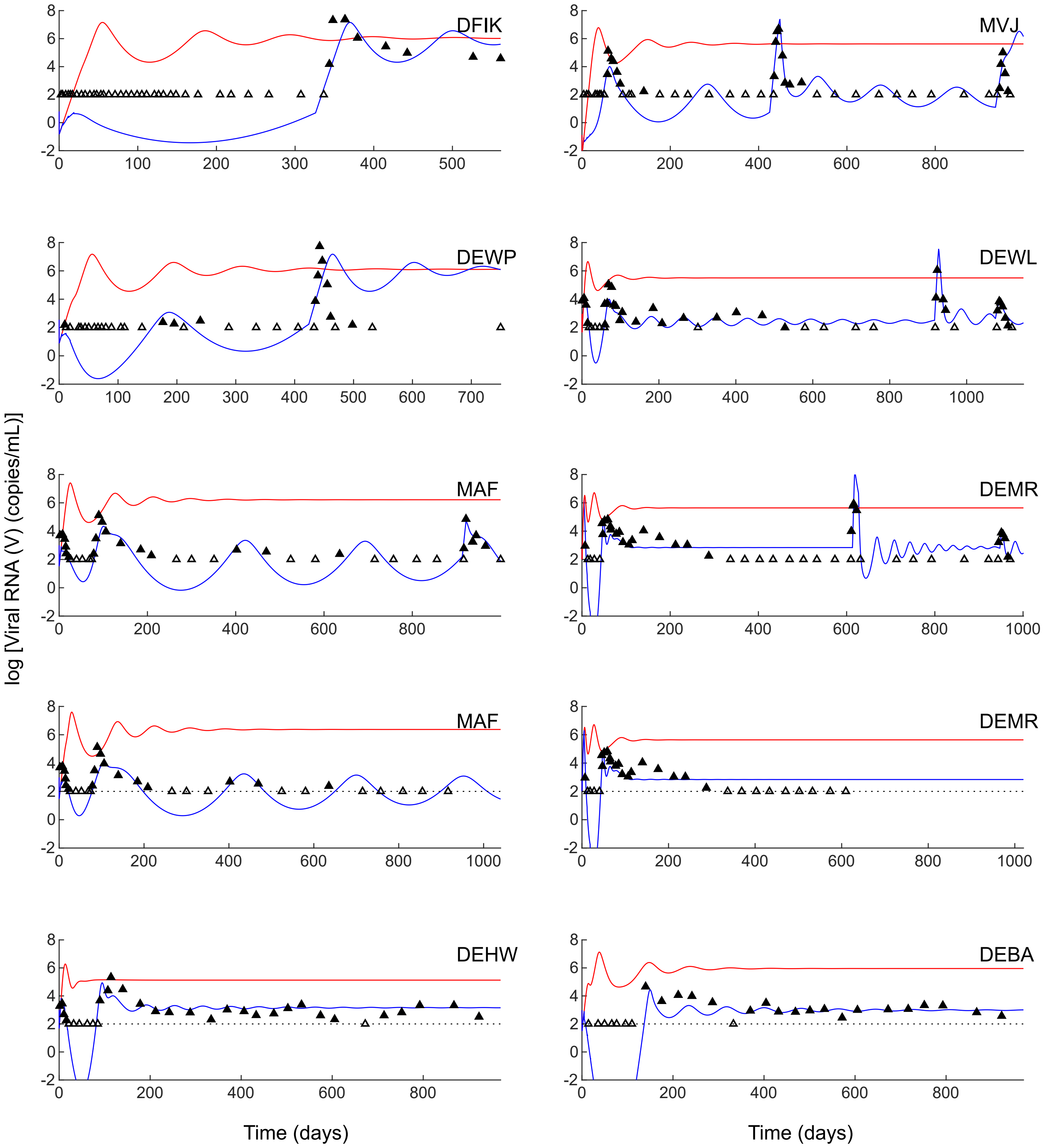


**Fig. S6 Hill coefficient,** $n = 3$. Fitting our model (Eqs. 11-18) with a Hill coefficient $n = 3$ following the procedure outlined in Methods yielded poor fits (blue) to the data (parameters in Table S7; also see comments in Table 3). Corresponding predictions without treatment are in red.

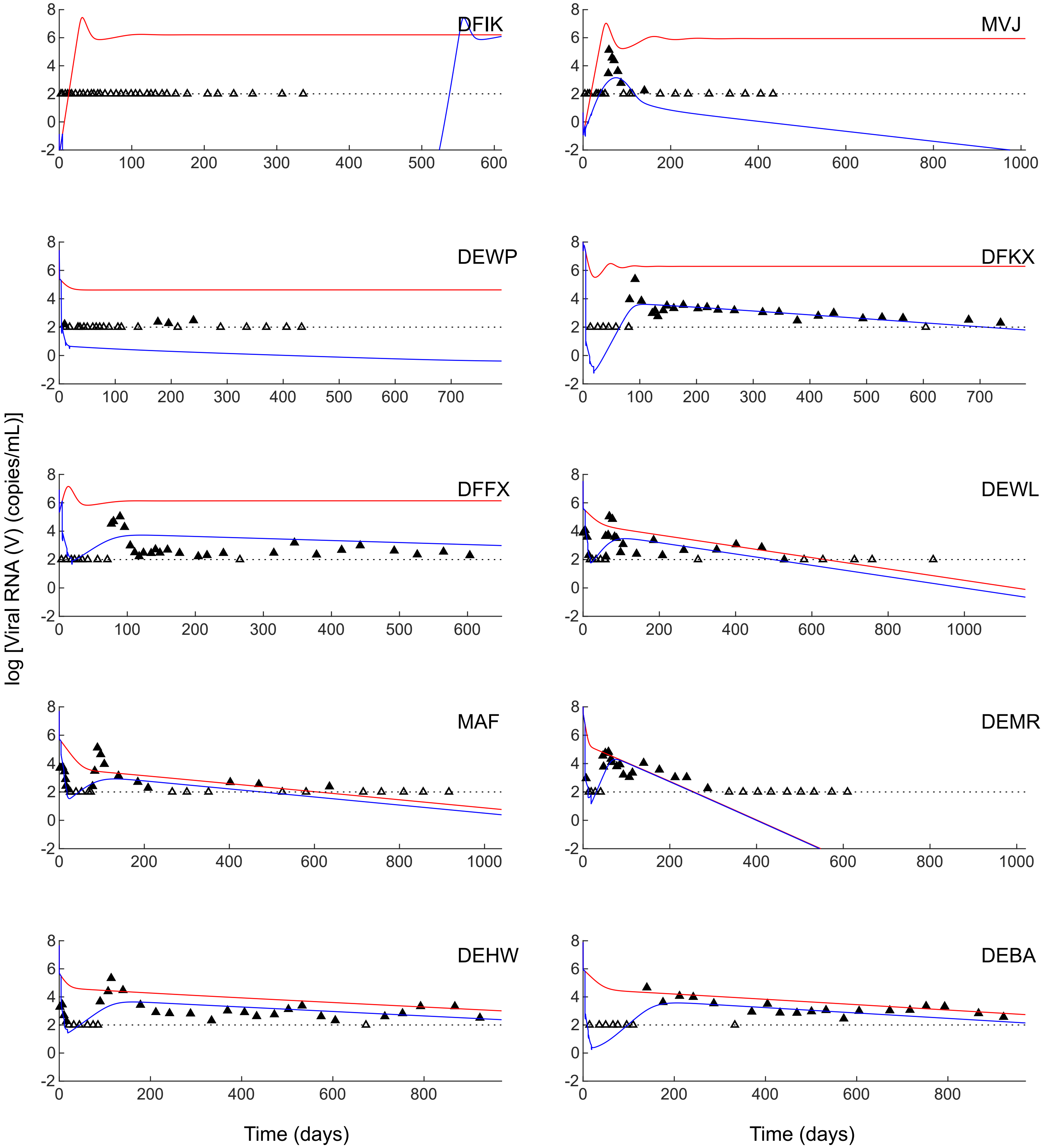


**Fig. S7 Absence of effector response.** Fitting with a model without a effector response (Eqs. 30-33) following the procedure outlined in Methods yielded poor fits (blue) to the data (parameters in Table S8). Corresponding predictions without treatment are in red.

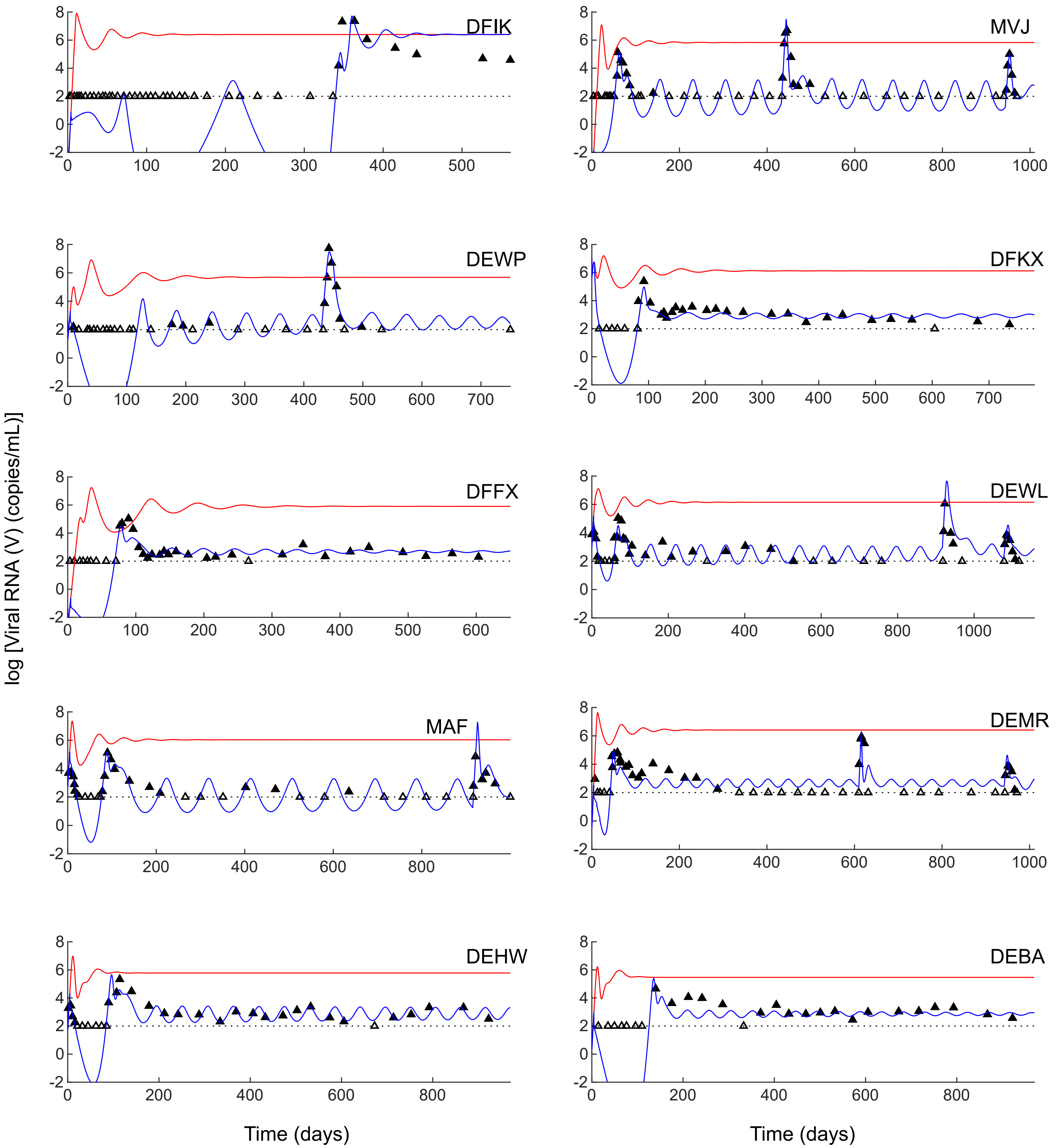


**Fig. S8 Varying effector proliferation ($k_E$).** Fitting with a model with varying effector proliferation ($k_E$ in Eq. 14) following the procedure outlined in Methods yielded good fits (blue) to the data (parameters in Table S9) but with a higher AIC (Table 3). Corresponding predictions without treatment are in red.

**Table S1** Individual parameter estimates for treated macaques obtained by simultaneously fitting our model (Eqs. 11-18) to $V$, $A_1$ and $A_2$ across both untreated macaques and responders (see Methods, Figures 2-4). Parameters pertaining to effector depletion experiments with anti-CD8$\alpha$ and anti-CD8$\beta$ antibodies were obtained from individual fits (best fits in Figure 2). The units of all the parameters are the same as in Table 2; $\zeta_\alpha$ and $\zeta_\beta$ are dimensionless while $\theta_m$, $\theta_\alpha$, $\theta_\beta$ are in days. (NA - not applicable)

| | DEMR | MVJ | DEWP | DEWL | MAF | DFIK | DFKX | DFFX | DEHW | DEBA |
|---|---|---|---|---|---|---|---|---|---|---|
| $V(0)$ | 506.42 | 0.01 | 2.35 | 969.05 | 665.59 | 0.12 | 205.86 | 0.16 | 1311.13 | 11.48 |
| $\omega_1$ | 2.23 | 2.59 | 2.07 | 2.66 | 2.27 | 2.09 | 1.73 | 2.10 | 2.49 | 2.49 |
| $\omega_2$ | 1.00 | 1.09 | 1.02 | 1.20 | 1.06 | 1.24 | 1.10 | 1.22 | 1.06 | 0.98 |
| $\eta_1$ | 0.26 | 0.11 | 0.04 | 0.09 | 0.06 | 0.13 | 0.07 | 0.08 | 0.06 | 0.06 |
| $\eta_2$ | 0.23 | 0.28 | 0.04 | 0.09 | 0.07 | 0.07 | 0.12 | 0.20 | 0.07 | 0.05 |
| $Vol_1$ | 74.90 | 138.86 | 336.77 | 1187.04 | 1864.01 | 52.46 | 521.95 | 260.89 | 433.74 | 262.41 |
| $Vol_2$ | 539.42 | 695.70 | 1271.71 | 347.62 | 464.55 | 695.10 | 1292.10 | 433.17 | 972.15 | 1671.88 |
| $k_1$ | 3.66 | 97.42 | 80.20 | 2.63 | 139.42 | 58.99 | 55.13 | 70.54 | 91.09 | 433.80 |
| $k_2$ | 694.24 | 2509.54 | 46.01 | 202.07 | 66.98 | 5.28 | 1494.26 | 1096.20 | 344.08 | 41.67 |
| $K$ | 59.19 | 148.67 | 144.49 | 308.05 | 141.89 | 146.72 | 32.49 | 69.74 | 77.64 | 35.08 |
| $\beta$ | $1.07\times10^{-8}$ | $1.04\times10^{-8}$ | $1.08\times10^{-8}$ | $1.15\times10^{-8}$ | $1.02\times10^{-8}$ | $1.05\times10^{-8}$ | $1.15\times10^{-8}$ | $9.64\times10^{-9}$ | $1.01\times10^{-8}$ | $9.85\times10^{-9}$ |
| $p^*$ | $4.95\times10^{9}$ | $7.15\times10^{9}$ | $6.92\times10^{9}$ | $6.39\times10^{9}$ | $6.61\times10^{9}$ | $5.95\times10^{9}$ | $5.53\times10^{9}$ | $5.76\times10^{9}$ | $6.41\times10^{9}$ | $7.62\times10^{9}$ |
| $m^*$ | 10.58 | 10.09 | 10.48 | 11.13 | 12.08 | 10.90 | 11.45 | 10.07 | 10.34 | 11.28 |
| $d_E$ | $8.36\times10^{-3}$ | $6.47\times10^{-3}$ | $1.24\times10^{-2}$ | $1.48\times10^{-2}$ | $5.29\times10^{-3}$ | $6.73\times10^{-3}$ | $1.22\times10^{-2}$ | $1.26\times10^{-2}$ | $1.23\times10^{-2}$ | $2.03\times10^{-2}$ |
| $\phi^*$ | $5.02\times10^{-5}$ | $2.31\times10^{-5}$ | $4.15\times10^{-6}$ | $5.45\times10^{-5}$ | $4.55\times10^{-5}$ | $1.46\times10^{-6}$ | $1.18\times10^{-4}$ | $4.52\times10^{-5}$ | $9.09\times10^{-5}$ | $8.84\times10^{-5}$ |
| $\xi$ | 0.17 | 0.21 | 0.40 | 0.22 | 0.20 | 0.24 | 0.21 | 0.21 | 0.21 | 0.25 |
| $f^*$ | $3.36\times10^{-11}$ | $5.49\times10^{-8}$ | $5.08\times10^{-9}$ | $7.01\times10^{-11}$ | $5.45\times10^{-10}$ | $2.91\times10^{-9}$ | $1.82\times10^{-9}$ | $9.69\times10^{-10}$ | $3.07\times10^{-10}$ | $1.37\times10^{-10}$ |
| $\zeta_\alpha$ | 0.33 | 0.84 | 0.99 | 0.40 | 0.35 | 0.90 | NA | NA | NA | NA |
| $\zeta_\beta$ | 0.56 | 0.39 | NA | 0.46 | NA | NA | NA | NA | NA | NA |
| $\theta_m$ | 11.86 | 10.68 | 18.17 | 5.33 | 4.73 | 3.53 | NA | NA | NA | NA |
| $\theta_\alpha$ | 607.26 | 436.54 | 431.86 | 918.09 | 916.21 | 333.23 | NA | NA | NA | NA |
| $\theta_\beta$ | 942.50 | 939.18 | NA | 1072.86 | NA | NA | NA | NA | NA | NA |

**Table S2** Individual parameter estimates for the ten untreated macaques obtained by simultaneously fitting our model (Eqs. 11-18) to $V$, $A_1$ and $A_2$ across both untreated macaques and responders (see Methods and Figure 4).

| Macaque | UT1 | UT2 | UT3 | UT4 | UT5 | UT6 | UT7 | UT8 | UT9 | UT10 |
|---|---|---|---|---|---|---|---|---|---|---|
| $V(0)$ | 52.27 | 51.93 | 17.58 | 3.67 | 3.22 | 15.05 | 1076.07 | 393.87 | 243.44 | 681.12 |
| $\beta$ | $1.06\times10^{-8}$ | $1.13\times10^{-8}$ | $1.06\times10^{-8}$ | $9.61\times10^{-9}$ | $9.68\times10^{-9}$ | $1.22\times10^{-8}$ | $9.74\times10^{-9}$ | $1.11\times10^{-8}$ | $1.17\times10^{-8}$ | $8.86\times10^{-9}$ |
| $p^*$ | $7.83\times10^{9}$ | $6.55\times10^{9}$ | $7.23\times10^{9}$ | $7.85\times10^{9}$ | $7.49\times10^{9}$ | $5.12\times10^{9}$ | $8.56\times10^{9}$ | $7.20\times10^{9}$ | $7.83\times10^{9}$ | $8.07\times10^{9}$ |
| $m^*$ | 11.30 | 10.66 | 10.07 | 10.79 | 11.83 | 10.04 | 10.39 | 12.06 | 10.87 | 11.34 |
| $d_E$ | $8.63\times10^{-3}$ | $6.93\times10^{-3}$ | $7.13\times10^{-3}$ | $7.79\times10^{-3}$ | $6.57\times10^{-3}$ | $8.28\times10^{-3}$ | $7.60\times10^{-3}$ | $1.23\times10^{-2}$ | $4.78\times10^{-3}$ | $1.06\times10^{-2}$ |
| $\phi^*$ | $1.51\times10^{-5}$ | $5.44\times10^{-6}$ | $1.17\times10^{-4}$ | $3.57\times10^{-5}$ | $5.85\times10^{-5}$ | $1.98\times10^{-4}$ | $1.08\times10^{-5}$ | $4.49\times10^{-5}$ | $2.83\times10^{-5}$ | $5.11\times10^{-5}$ |
| $\xi$ | 0.32 | 0.18 | 0.54 | 0.49 | 0.29 | 0.43 | 0.41 | 0.56 | 0.33 | 0.18 |

**Table S3** Individual parameter estimates for treated macaques obtained by simultaneously fitting models without enhanced antigen clearance by bNAbs (no $AV$ term in Eq. 13) to $V$, $A_1$ and $A_2$ across both untreated macaques and responders (see Methods for details).

| | DEMR | MVJ | DEWP | DEWL | MAF | DFIK | DFKX | DFFX | DEHW | DEBA |
|---|---|---|---|---|---|---|---|---|---|---|
| $V(0)$ | 506.42 | 0.01 | 2.35 | 969.05 | 665.59 | 0.12 | 205.86 | 0.16 | 1311.13 | 11.48 |
| $\omega_1$ | 2.23 | 2.59 | 2.07 | 2.66 | 2.27 | 2.09 | 1.73 | 2.10 | 2.49 | 2.49 |
| $\omega_2$ | 1.00 | 1.09 | 1.02 | 1.20 | 1.06 | 1.24 | 1.10 | 1.22 | 1.06 | 0.98 |
| $\eta_1$ | 0.26 | 0.11 | 0.04 | 0.09 | 0.06 | 0.13 | 0.07 | 0.08 | 0.06 | 0.06 |
| $\eta_2$ | 0.23 | 0.28 | 0.04 | 0.09 | 0.07 | 0.07 | 0.12 | 0.20 | 0.07 | 0.05 |
| $Vol_1$ | 74.90 | 138.86 | 336.77 | 1187.04 | 1864.01 | 52.46 | 521.95 | 260.89 | 433.74 | 262.41 |
| $Vol_2$ | 539.42 | 695.70 | 1271.71 | 347.62 | 464.55 | 695.10 | 1292.10 | 433.17 | 972.15 | 1671.88 |
| $k_1$ | 3.66 | 97.42 | 80.20 | 2.63 | 139.42 | 58.99 | 55.13 | 70.54 | 91.09 | 433.80 |
| $k_2$ | 694.24 | 2509.54 | 46.01 | 202.07 | 66.98 | 5.28 | 1494.26 | 1096.20 | 344.08 | 41.67 |
| $K$ | 59.19 | 148.67 | 144.49 | 308.05 | 141.89 | 146.72 | 32.49 | 69.74 | 77.64 | 35.08 |
| $\beta$ | $1.07\times10^{-8}$ | $1.04\times10^{-8}$ | $1.08\times10^{-8}$ | $1.15\times10^{-8}$ | $1.02\times10^{-8}$ | $1.05\times10^{-8}$ | $1.15\times10^{-8}$ | $9.64\times10^{-9}$ | $1.01\times10^{-8}$ | $9.85\times10^{-9}$ |
| $p^*$ | $4.95\times10^{9}$ | $7.15\times10^{9}$ | $6.92\times10^{9}$ | $6.39\times10^{9}$ | $6.61\times10^{9}$ | $5.95\times10^{9}$ | $5.53\times10^{9}$ | $5.76\times10^{9}$ | $6.41\times10^{9}$ | $7.62\times10^{9}$ |
| $m^*$ | 10.58 | 10.09 | 10.48 | 11.13 | 12.08 | 10.90 | 11.45 | 10.07 | 10.34 | 11.28 |
| $d_E$ | $8.36\times10^{-3}$ | $6.47\times10^{-3}$ | $1.24\times10^{-2}$ | $1.48\times10^{-2}$ | $5.29\times10^{-3}$ | $6.73\times10^{-3}$ | $1.22\times10^{-2}$ | $1.26\times10^{-2}$ | $1.23\times10^{-2}$ | $2.03\times10^{-2}$ |
| $\phi^*$ | $5.02\times10^{-5}$ | $2.31\times10^{-5}$ | $4.15\times10^{-6}$ | $5.45\times10^{-5}$ | $4.55\times10^{-5}$ | $1.46\times10^{-6}$ | $1.18\times10^{-4}$ | $4.52\times10^{-5}$ | $9.09\times10^{-5}$ | $8.84\times10^{-5}$ |
| $\xi$ | 0.17 | 0.21 | 0.40 | 0.22 | 0.20 | 0.24 | 0.21 | 0.21 | 0.21 | 0.25 |
| $f^*$ | $3.36\times10^{-11}$ | $5.49\times10^{-8}$ | $5.08\times10^{-9}$ | $7.01\times10^{-11}$ | $5.45\times10^{-10}$ | $2.91\times10^{-9}$ | $1.82\times10^{-9}$ | $9.69\times10^{-10}$ | $3.07\times10^{-10}$ | $1.37\times10^{-10}$ |
| $\zeta_\alpha$ | 0.33 | 0.84 | 0.99 | 0.40 | 0.35 | 0.90 | NA | NA | NA | NA |
| $\zeta_\beta$ | 0.56 | 0.39 | NA | 0.46 | NA | NA | NA | NA | NA | NA |
| $\theta_m$ | 11.86 | 10.68 | 18.17 | 5.33 | 4.73 | 3.53 | NA | NA | NA | NA |
| $\theta_\alpha$ | 607.26 | 436.54 | 431.86 | 918.09 | 916.21 | 333.23 | NA | NA | NA | NA |
| $\theta_\beta$ | 942.50 | 939.18 | NA | 1072.86 | NA | NA | NA | NA | NA | NA |

**Table S4** Individual parameter estimates for treated macaques obtained by simultaneously fitting models without enhanced antigen uptake and subsequent effector elicitation (no $f^*AV$ term in Eq. 14) by bNAbs to $V$, $A_1$ and $A_2$ across both untreated macaques and responders (see Methods for details).

| | DEMR | MVJ | DEWP | DEWL | MAF | DFIK | DFKX | DFFX | DEHW | DEBA |
|---|---|---|---|---|---|---|---|---|---|---|
| $V(0)$ | 8.76E+04 | 4.52E-05 | 6.20E-01 | 1.72E+06 | 2.35E+05 | 4.42E+00 | 6.81E+03 | 4.39E-03 | 3.48E+03 | 1.53E-03 |
| $\omega_1$ | 1.51 | 1.71 | 2.54 | 2.57 | 2.09 | 2.33 | 2.16 | 2.02 | 2.33 | 1.78 |
| $\omega_2$ | 0.99 | 1.26 | 1.17 | 1.17 | 1.35 | 1.46 | 1.05 | 1.09 | 1.12 | 1.20 |
| $\eta_1$ | 0.24 | 0.12 | 0.04 | 0.07 | 0.07 | 0.11 | 0.07 | 0.11 | 0.07 | 0.06 |
| $\eta_2$ | 0.18 | 0.26 | 0.04 | 0.09 | 0.08 | 0.07 | 0.23 | 0.23 | 0.07 | 0.06 |
| $Vol_1$ | 76.79 | 90.56 | 458.29 | 2668.51 | 1590.33 | 90.40 | 686.27 | 128.48 | 433.49 | 252.96 |
| $Vol_2$ | 529.42 | 818.14 | 1006.27 | 372.44 | 441.28 | 854.29 | 736.39 | 265.97 | 1011.31 | 1349.65 |
| $k_1$ | 1.48E-09 | 6.41E-18 | 2.91E-02 | 5.58E-12 | 3.90E-05 | 5.27E-10 | 5.29E-05 | 2.94E-05 | 8.12E-09 | 1.02E+01 |
| $k_2$ | 13.37 | 34.20 | 88.56 | 30.71 | 34.26 | 68.47 | 13.95 | 106.02 | 57.79 | 54.87 |
| $K$ | 1116.77 | 382.85 | 5047.78 | 22845.91 | 58520.72 | 36852.02 | 546.43 | 984.93 | 5159.19 | 517.12 |
| $\beta$ | 3.28E-08 | 1.69E-08 | 6.28E-09 | 6.06E-09 | 9.95E-09 | 1.29E-08 | 1.37E-08 | 1.44E-08 | 8.94E-09 | 7.55E-09 |
| $p^*$ | 1.49E+10 | 2.08E+09 | 6.23E+09 | 5.64E+09 | 2.98E+09 | 2.39E+09 | 3.99E+09 | 8.34E+09 | 3.89E+09 | 4.82E+09 |
| $m^*$ | 310.52 | 1.56 | 12.47 | 8.61 | 5.18 | 13.17 | 18.64 | 34.81 | 6.22 | 12.38 |
| $d_E$ | 8.96E-04 | 6.33E-03 | 1.61E-02 | 5.31E-02 | 4.81E-02 | 5.29E-03 | 3.78E-02 | 3.50E-02 | 3.05E-02 | 1.66E-02 |
| $\phi^*$ | 5.36E-05 | 1.52E-06 | 7.74E-06 | 1.91E-04 | 2.50E-05 | 6.41E-06 | 1.41E-04 | 3.65E-05 | 1.28E-04 | 3.38E-04 |
| $\xi$ | 0.30 | 0.10 | 7.91 | 9.74 | 4.92 | 6.14 | 10.32 | 3.80 | 5.46 | 28.05 |
| $f^*$ | 7.19E-07 | 1.42E-06 | 2.95E-07 | 8.14E-07 | 1.24E-06 | 5.88E-06 | 2.37E-07 | 8.33E-06 | 2.22E-07 | 3.35E-07 |

**Table S5** Individual parameter estimates obtained by fitting a model with an alternative description of effector exhaustion (Eqs. 24-28; see Methods for details)

| | DFIK | MVJ | DEWP | DFKX | DFFX | DEWL | MAF | DEMR | DEHW | DEBA |
|---|---|---|---|---|---|---|---|---|---|---|
| $V(0)$ | $9.86\times10^{2}$ | $1.98\times10^{3}$ | $1.66\times10^{2}$ | $2.94\times10^{2}$ | $1.53\times10^{-2}$ | $6.01\times10^{4}$ | $3.57\times10^{3}$ | $2.47\times10^{3}$ | $1.61\times10^{4}$ | $1.42\times10^{3}$ |
| $\omega_1$ | 1.71 | 2.09 | 1.10 | 1.51 | 1.60 | 2.55 | 1.36 | 1.58 | 1.65 | 1.72 |
| $\omega_2$ | 1.14 | 1.32 | 1.12 | 1.48 | 0.85 | 0.86 | 0.84 | 1.17 | 1.74 | 1.19 |
| $\eta_1$ | 0.12 | 0.10 | 0.04 | 0.07 | 0.10 | 0.08 | 0.08 | 0.26 | 0.07 | 0.06 |
| $\eta_2$ | 0.07 | 0.23 | 0.04 | 0.25 | 0.16 | 0.10 | 0.07 | 0.18 | 0.07 | 0.07 |
| $Vol_1$ | 58.22 | 156.47 | 465.69 | 618.50 | 220.32 | 1644.68 | 1201.31 | 53.05 | 323.90 | 276.45 |
| $Vol_2$ | 764.22 | 901.37 | 1250.82 | 575.51 | 481.40 | 258.48 | 418.49 | 782.81 | 1066.82 | 1272.74 |
| $k_1$ | 0.00 | 0.00 | 2510.94 | 0.00 | 0.00 | 0.00 | 30708.34 | 0.00 | 0.00 | 0.00 |
| $k_2$ | 1588.14 | 294.24 | 162.13 | 1792.36 | 383.87 | 232.45 | 111.35 | 181.36 | 392.91 | 291.91 |
| $K$ | 457.27 | 798.00 | 139.83 | 250.77 | 2241.92 | 1516.07 | 1354.24 | 1359.91 | 133.12 | 81.76 |
| $\beta$ | $1.04\times10^{-8}$ | $1.94\times10^{-8}$ | $5.53\times10^{-8}$ | $1.90\times10^{-8}$ | $1.87\times10^{-8}$ | $2.00\times10^{-8}$ | $4.85\times10^{-8}$ | $1.23\times10^{-8}$ | $2.03\times10^{-8}$ | $1.22\times10^{-8}$ |
| $p^*$ | $2.53\times10^{9}$ | $1.87\times10^{9}$ | $2.35\times10^{9}$ | $2.12\times10^{9}$ | $2.17\times10^{9}$ | $1.48\times10^{9}$ | $2.19\times10^{9}$ | $2.27\times10^{9}$ | $1.55\times10^{9}$ | $2.75\times10^{9}$ |
| $m^*$ | 0.15 | 12.71 | 0.56 | 0.23 | 2.37 | 0.06 | 0.22 | 0.28 | 1.25 | 0.21 |
| $d_E$ | $1.35\times10^{-2}$ | $2.44\times10^{-3}$ | $1.07\times10^{-3}$ | $9.04\times10^{-2}$ | $1.12\times10^{-2}$ | $7.22\times10^{-2}$ | $2.01\times10^{-2}$ | $2.02\times10^{-2}$ | $4.16\times10^{-5}$ | $4.25\times10^{-2}$ |
| $\phi_1^*$ | $2.27\times10^{-1}$ | $5.27\times10^{-3}$ | $5.53\times10^{-7}$ | $2.05\times10^{-5}$ | $4.21\times10^{-6}$ | $5.32\times10^{-6}$ | $9.18\times10^{-6}$ | $9.94\times10^{-5}$ | $7.58\times10^{-6}$ | $3.79\times10^{-5}$ |
| $\xi$ | 0.07 | 0.07 | 0.07 | 0.07 | 0.12 | 0.10 | 0.11 | 0.08 | 0.07 | 0.10 |
| $\phi_2^*$ | $7.48\times10^{-6}$ | $5.30\times10^{-7}$ | $3.95\times10^{-6}$ | $1.55\times10^{-3}$ | $4.62\times10^{-11}$ | $3.44\times10^{-4}$ | $1.27\times10^{-4}$ | $6.07\times10^{-5}$ | $8.31\times10^{-7}$ | $1.08\times10^{-4}$ |
| $f^*$ | 0 | 0 | 0 | 0 | 0 | 0 | 0 | 0 | 0 | 0 |

**Table S6** Individual parameter estimates obtained as in Table S1 but with the Hill coefficient $n = 1$ (see Methods for details).

| | DFIK | MVJ | DEWP | DFKX | DFFX | DEWL | MAF | DEMR | DEHW | DEBA |
|---|---|---|---|---|---|---|---|---|---|---|
| $V(0)$ | $7.84\times10^{-3}$ | $9.42\times10^{-3}$ | $1.58\times10^{3}$ | $2.67\times10^{-1}$ | $1.42\times10^{-1}$ | $4.67\times10^{4}$ | $1.68\times10^{5}$ | $7.01\times10^{2}$ | $3.99\times10^{3}$ | $3.00\times10^{0}$ |
| $\omega_1$ | 1.31 | 2.21 | 0.97 | 2.00 | 1.54 | 1.43 | 0.85 | 1.61 | 2.27 | 1.61 |
| $\omega_2$ | 0.89 | 1.67 | 1.04 | 2.43 | 2.32 | 1.58 | 1.08 | 2.05 | 1.48 | 1.43 |
| $\eta_1$ | 0.12 | 0.10 | 0.04 | 0.06 | 0.10 | 0.07 | 0.07 | 0.28 | 0.07 | 0.06 |
| $\eta_2$ | 0.07 | 0.25 | 0.05 | 0.25 | 0.14 | 0.09 | 0.07 | 0.27 | 0.07 | 0.06 |
| $Vol_1$ | 64.72 | 138.60 | 502.26 | 627.41 | 136.64 | 1783.29 | 1487.30 | 47.12 | 400.49 | 247.16 |
| $Vol_2$ | 641.78 | 834.52 | 1088.40 | 683.72 | 637.74 | 317.29 | 555.65 | 479.13 | 1036.00 | 1444.39 |
| $k_1$ | 0.81 | 0.02 | 0.35 | 0.70 | 0.28 | 2.92 | 0.30 | 0.29 | 0.47 | 0.13 |
| $k_2$ | 56.94 | 3027.28 | 4.21 | 138.79 | 55.87 | 39.10 | 50.87 | 65.15 | 41766.23 | 1047.63 |
| $K$ | $1.85\times10^{-5}$ | $6.07\times10^{-8}$ | $2.29\times10^{-3}$ | $7.54\times10^{-4}$ | 0.04 | 162.74 | 153.32 | 35.28 | 2364.55 | 36.66 |
| $\beta$ | $8.72\times10^{-9}$ | $8.46\times10^{-9}$ | $7.23\times10^{-9}$ | $9.60\times10^{-9}$ | $6.89\times10^{-9}$ | $1.80\times10^{-8}$ | $7.61\times10^{-9}$ | $5.39\times10^{-9}$ | $3.10\times10^{-8}$ | $6.00\times10^{-9}$ |
| $p^*$ | $4.65\times10^{9}$ | $6.94\times10^{9}$ | $3.13\times10^{9}$ | $3.04\times10^{9}$ | $4.26\times10^{9}$ | $3.44\times10^{9}$ | $5.88\times10^{9}$ | $5.20\times10^{9}$ | $2.74\times10^{9}$ | $8.55\times10^{9}$ |
| $m^*$ | 2.73 | 41.58 | 1.81 | 2.71 | 6.78 | 483.58 | 5180.91 | 26.52 | 9.59 | 0.30 |
| $d_E$ | $3.82\times10^{-2}$ | $7.74\times10^{-4}$ | $2.33\times10^{-2}$ | $4.21\times10^{-1}$ | $3.51\times10^{-3}$ | $8.25\times10^{-4}$ | $2.81\times10^{-4}$ | $1.63\times10^{-5}$ | $2.99\times10^{-5}$ | $6.49\times10^{-4}$ |
| $\phi^*$ | $3.11\times10^{-3}$ | $1.62\times10^{-3}$ | $3.06\times10^{-3}$ | $1.44\times10^{-4}$ | $6.59\times10^{-5}$ | $5.81\times10^{-4}$ | $6.29\times10^{-3}$ | $3.72\times10^{-3}$ | $1.01\times10^{-4}$ | $2.34\times10^{-5}$ |
| $\xi$ | 0.65 | 0.32 | 0.02 | 0.17 | 0.09 | 0.19 | 0.80 | 0.04 | 0.06 | 0.07 |
| $f^*$ | $1.93\times10^{-4}$ | $2.12\times10^{-5}$ | $1.38\times10^{-5}$ | $2.67\times10^{-9}$ | $4.15\times10^{-9}$ | $3.50\times10^{-13}$ | $1.24\times10^{-10}$ | $8.23\times10^{-6}$ | $1.92\times10^{-11}$ | $7.18\times10^{-8}$ |

**Table S7** Individual parameter estimates obtained as in Table S1 but with the Hill coefficient $n = 3$ (see Methods for details).

| | DFIK | MVJ | DEWP | DFKX | DFFX | DEWL | MAF | DEMR | DEHW | DEBA |
|---|---|---|---|---|---|---|---|---|---|---|
| $V(0)$ | $6.34\times10^{0}$ | $1.33\times10^{-1}$ | $3.53\times10^{2}$ | $9.87\times10^{2}$ | $1.00\times10^{3}$ | $9.28\times10^{2}$ | $9.99\times10^{2}$ | $9.99\times10^{2}$ | $1.00\times10^{3}$ | $9.53\times10^{2}$ |
| $\omega_1$ | 1.78 | 1.27 | 1.58 | 1.32 | 1.89 | 1.30 | 1.80 | 1.85 | 1.59 | 1.72 |
| $\omega_2$ | 1.78 | 1.81 | 1.77 | 1.88 | 1.96 | 1.75 | 2.31 | 1.90 | 2.44 | 1.97 |
| $\eta_1$ | 0.12 | 0.13 | 0.04 | 0.07 | 0.10 | 0.08 | 0.06 | 0.21 | 0.07 | 0.06 |
| $\eta_2$ | 0.08 | 0.15 | 0.05 | 0.18 | 0.17 | 0.09 | 0.07 | 0.14 | 0.07 | 0.07 |
| $Vol_1$ | 80.20 | 80.30 | 548.87 | 608.11 | 167.20 | 1630.38 | 2194.97 | 117.92 | 387.75 | 243.09 |
| $Vol_2$ | 601.75 | 1470.23 | 1037.82 | 866.97 | 564.90 | 369.15 | 477.42 | 832.63 | 1222.64 | 1501.78 |
| $k_1$ | 54.91 | 33.41 | 21.63 | 75.68 | 24.69 | 47.39 | 46.89 | 59.66 | 50.04 | 58.85 |
| $k_2$ | 87.54 | 377.98 | 162.26 | 177.43 | 1349.73 | 377.25 | 64.73 | 512.79 | 303.04 | 505.15 |
| $K$ | 3067.59 | 104.77 | 570.35 | 158.55 | 488.02 | 1468.32 | 28.57 | 453.75 | 141.67 | 56.91 |
| $\beta$ | $6.53\times10^{-9}$ | $1.13\times10^{-8}$ | $4.68\times10^{-9}$ | $1.11\times10^{-8}$ | $6.45\times10^{-9}$ | $1.47\times10^{-8}$ | $5.38\times10^{-9}$ | $1.44\times10^{-8}$ | $1.47\times10^{-8}$ | $8.28\times10^{-9}$ |
| $p^*$ | $5.20\times10^{9}$ | $4.15\times10^{9}$ | $6.82\times10^{9}$ | $4.52\times10^{9}$ | $5.79\times10^{9}$ | $4.45\times10^{9}$ | $8.52\times10^{9}$ | $8.91\times10^{9}$ | $4.32\times10^{9}$ | $5.61\times10^{9}$ |
| $m^*$ | 16.26 | 15.01 | 28.81 | 17.39 | 25.29 | 17.42 | 13.23 | 36.92 | 13.29 | 29.18 |
| $d_E$ | $1.06\times10^{-3}$ | $1.97\times10^{-3}$ | $2.74\times10^{-3}$ | $8.47\times10^{-3}$ | $7.99\times10^{-3}$ | $5.30\times10^{-3}$ | $1.44\times10^{-3}$ | $1.64\times10^{-2}$ | $9.84\times10^{-3}$ | $8.79\times10^{-3}$ |
| $\phi^*$ | $2.65\times10^{-5}$ | $5.78\times10^{-5}$ | $7.59\times10^{-5}$ | $1.96\times10^{-4}$ | $1.97\times10^{-4}$ | $7.21\times10^{-5}$ | $1.05\times10^{-4}$ | $5.06\times10^{-5}$ | $2.58\times10^{-4}$ | $3.29\times10^{-4}$ |
| $\xi$ | 1.08 | 0.21 | 3.92 | 1.28 | 1.64 | 0.26 | 0.26 | 0.52 | 0.24 | 1.51 |
| $f^*$ | $5.28\times10^{-7}$ | $3.22\times10^{-8}$ | $3.67\times10^{-8}$ | $6.85\times10^{-9}$ | $5.92\times10^{-9}$ | $1.17\times10^{-9}$ | $3.69\times10^{-9}$ | $3.29\times10^{-10}$ | $9.08\times10^{-10}$ | $9.36\times10^{-9}$ |
| $\zeta_\alpha$ | 0.99 | 0.00 | 0.38 | NA | NA | 1.00 | 0.00 | 1.00 | NA | NA |
| $\zeta_\beta$ | NA | 0.67 | NA | NA | NA | 0.44 | NA | 0.34 | NA | NA |
| $\theta_m$ | 30.00 | 23.16 | 25.06 | NA | NA | 10.74 | 8.36 | 13.69 | NA | NA |
| $\theta_\alpha$ | 326.00 | 425.05 | 424.00 | NA | NA | 916.51 | 913.15 | 613.24 | NA | NA |
| $\theta_\beta$ | NA | 936.50 | NA | NA | NA | 1074.76 | NA | 947.50 | NA | NA |

**Table S8** Individual parameter estimates obtained by fitting a model without an explicit effector response (Eqs. 30-33; see Methods for details). Here, $d_L$ was fixed at $0.004$ day$^{-1}$.

| | DFIK | MVJ | DEWP | DFKX | DFFX | DEWL | MAF | DEMR | DEHW | DEBA |
|---|---|---|---|---|---|---|---|---|---|---|
| $V(0)$ | $1.36\times10^{-1}$ | $4.75\times10^{0}$ | $2.68\times10^{7}$ | $3.22\times10^{8}$ | $8.40\times10^{6}$ | $3.44\times10^{7}$ | $5.11\times10^{7}$ | $7.17\times10^{9}$ | $4.77\times10^{7}$ | $7.89\times10^{7}$ |
| $\omega_1$ | 2.03 | 2.23 | 1.63 | 2.11 | 2.06 | 1.88 | 1.76 | 2.54 | 1.52 | 2.02 |
| $\omega_2$ | 1.50 | 1.12 | 1.02 | 1.69 | 1.36 | 2.47 | 1.24 | 1.49 | 1.04 | 1.20 |
| $\eta_1$ | 0.12 | 0.11 | 0.04 | 0.07 | 0.10 | 0.09 | 0.07 | 0.21 | 0.07 | 0.06 |
| $\eta_2$ | 0.08 | 0.08 | 0.05 | 0.24 | 0.19 | 0.10 | 0.06 | 0.20 | 0.07 | 0.08 |
| $Vol_1$ | 77.57 | 119.51 | 411.73 | 666.02 | 199.69 | 1154.70 | 1231.05 | 126.27 | 402.65 | 179.70 |
| $Vol_2$ | 600.62 | 1425.75 | 977.06 | 534.87 | 435.31 | 354.16 | 535.88 | 541.10 | 1119.65 | 1328.18 |
| $k_1$ | $5.18\times10^{3}$ | $9.52\times10^{-1}$ | $5.97\times10^{-3}$ | $5.32\times10^{2}$ | $6.42\times10^{1}$ | $7.74\times10^{2}$ | $6.67\times10^{1}$ | $7.47\times10^{0}$ | $3.33\times10^{4}$ | $6.63\times10^{-1}$ |
| $k_2$ | $9.25\times10^{3}$ | $1.15\times10^{2}$ | $1.40\times10^{5}$ | $1.15\times10^{8}$ | $1.71\times10^{4}$ | $5.74\times10^{5}$ | 2.01E+13 | $7.38\times10^{8}$ | $6.97\times10^{4}$ | $3.90\times10^{5}$ |
| $K$ | $8.27\times10^{-12}$ | $1.00\times10^{-35}$ | $6.06\times10^{-52}$ | $3.67\times10^{-27}$ | $6.84\times10^{-38}$ | $1.68\times10^{5}$ | $8.43\times10^{12}$ | $4.04\times10^{5}$ | $2.17\times10^{3}$ | $2.35\times10^{2}$ |
| $\beta$ | $1.18\times10^{-8}$ | $9.84\times10^{-9}$ | $1.04\times10^{-8}$ | $2.46\times10^{-8}$ | $9.56\times10^{-9}$ | $6.06\times10^{-9}$ | $4.56\times10^{-9}$ | $5.39\times10^{-9}$ | $5.78\times10^{-9}$ | $3.62\times10^{-9}$ |
| $p^*$ | $4.78\times10^{9}$ | $3.67\times10^{9}$ | $2.08\times10^{9}$ | $4.80\times10^{9}$ | $5.47\times10^{9}$ | $2.94\times10^{9}$ | $3.59\times10^{9}$ | $2.27\times10^{9}$ | $3.26\times10^{9}$ | $5.12\times10^{9}$ |
| $f_L$ | $1.29\times10^{-1}$ | $6.37\times10^{-2}$ | $3.12\times10^{-1}$ | $1.24\times10^{-1}$ | $3.09\times10^{-1}$ | $5.33\times10^{-2}$ | $3.98\times10^{-2}$ | $6.18\times10^{-2}$ | $2.20\times10^{-1}$ | $1.04\times10^{-1}$ |
| $\psi$ | $2.87\times10^{-4}$ | $2.39\times10^{-4}$ | $2.32\times10^{-4}$ | $2.85\times10^{-4}$ | $3.35\times10^{-4}$ | $2.95\times10^{-4}$ | $6.02\times10^{-5}$ | $1.83\times10^{-3}$ | $3.61\times10^{-4}$ | $8.09\times10^{-5}$ |
| $a$ | $1.48\times10^{-2}$ | $4.68\times10^{-3}$ | $9.90\times10^{-3}$ | $3.02\times10^{-3}$ | $3.69\times10^{-3}$ | $1.21\times10^{-2}$ | $3.35\times10^{-3}$ | $3.34\times10^{-2}$ | $9.75\times10^{-3}$ | $3.89\times10^{-3}$ |

**Table S9** Individual parameter estimates obtained as in Table S1 but with varying effector proliferation rate, $k_E$ (see Methods for details).

| | DFIK | MVJ | DEWP | DFKX | DFFX | DEWL | MAF | DEMR | DEHW | DEBA |
|---|---|---|---|---|---|---|---|---|---|---|
| $V(0)$ | 5.29E+00 | 6.86E-04 | 4.45E+02 | 2.18E+04 | 1.05E+04 | 2.33E-03 | 9.24E+06 | 4.66E-02 | 1.13E+03 | 4.18E+01 |
| $\omega_1$ | 1.72 | 2.26 | 1.38 | 2.12 | 1.40 | 1.94 | 2.46 | 1.48 | 1.57 | 1.81 |
| $\omega_2$ | 1.27 | 1.57 | 1.45 | 1.72 | 1.46 | 1.13 | 1.94 | 1.43 | 1.52 | 1.48 |
| $\eta_1$ | 0.20 | 0.10 | 0.04 | 0.08 | 0.07 | 0.11 | 0.07 | 0.10 | 0.07 | 0.06 |
| $\eta_2$ | 0.25 | 0.11 | 0.05 | 0.09 | 0.08 | 0.07 | 0.20 | 0.15 | 0.08 | 0.06 |
| $Vol_1$ | 162.56 | 151.08 | 445.13 | 1699.25 | 1448.49 | 75.22 | 649.98 | 209.87 | 404.28 | 297.63 |
| $Vol_2$ | 766.44 | 1283.45 | 1112.86 | 338.39 | 444.12 | 760.72 | 890.94 | 625.08 | 876.11 | 1599.91 |
| $k_1$ | 225.35 | 89.51 | 263.41 | 190.01 | 240.82 | 186.99 | 235.32 | 156.06 | 214.53 | 354.58 |
| $k_2$ | 517.34 | 355.60 | 206.67 | 99.75 | 158.40 | 157.59 | 192.37 | 107.03 | 383.18 | 315.02 |
| $K$ | 96.71 | 113.75 | 130.45 | 65.22 | 56.66 | 46.37 | 59.92 | 54.12 | 57.23 | 25.85 |
| $\beta$ | 1.20E-08 | 1.74E-08 | 1.79E-08 | 1.26E-08 | 1.62E-08 | 2.21E-08 | 1.18E-08 | 1.59E-08 | 1.41E-08 | 1.34E-08 |
| $p^*$ | 7.15E+09 | 4.15E+09 | 3.45E+09 | 5.02E+09 | 4.44E+09 | 5.67E+09 | 5.49E+09 | 3.90E+09 | 5.10E+09 | 5.49E+09 |
| $m^*$ | 3.59 | 3.55 | 11.30 | 4.67 | 4.46 | 2.81 | 13.16 | 9.11 | 4.29 | 8.82 |
| $d_E$ | 1.17E-02 | 8.67E-03 | 1.50E-02 | 1.02E-02 | 5.14E-03 | 1.46E-03 | 9.75E-03 | 1.83E-02 | 1.10E-02 | 1.36E-02 |
| $k_E$ | 0.38 | 0.40 | 0.35 | 0.40 | 0.37 | 0.34 | 0.44 | 0.33 | 0.43 | 0.32 |
| $\phi^*$ | 8.76E-05 | 1.07E-04 | 1.24E-04 | 1.34E-04 | 2.33E-04 | 1.02E-04 | 3.16E-04 | 1.17E-04 | 2.75E-04 | 1.45E-04 |
| $\xi$ | 1.24 | 0.49 | 0.82 | 0.93 | 0.51 | 1.22 | 1.24 | 1.13 | 0.51 | 0.44 |
| $f^*$ | 7.71E-09 | 3.18E-09 | 6.96E-11 | 8.38E-11 | 1.48E-11 | 5.88E-08 | 3.82E-12 | 7.21E-10 | 1.65E-12 | 2.63E-11 |
| $\zeta_\alpha$ | 0.39 | 0.94 | 0.95 | 0.85 | 0.89 | 0.57 | NA | NA | NA | NA |
| $\zeta_\beta$ | 0.57 | 0.76 | NA | 0.75 | NA | NA | NA | NA | NA | NA |
| $\theta_m$ | 5.38 | 10.87 | 20.16 | 14.37 | 10.75 | 2.75 | NA | NA | NA | NA |
| $\theta_\alpha$ | 609.98 | 432.62 | 429.95 | 916.98 | 915.27 | 330.76 | NA | NA | NA | NA |
| $\theta_\beta$ | 942.11 | 943.54 | NA | 1077.44 | NA | NA | NA | NA | NA | NA |
